# Somatic genetic drift and multi-level selection in modular species

**DOI:** 10.1101/833335

**Authors:** Lei Yu, Christoffer Boström, Sören Franzenburg, Till Bayer, Tal Dagan, Thorsten B.H. Reusch

## Abstract

Cells in multicellular organisms are genetically heterogeneous owing to somatic mutations. The accumulation of somatic genetic variation in species undergoing asexual (or clonal) reproduction (termed modular species) may lead to phenotypic heterogeneity among modules. However, abundance and dynamics of somatic genetic variation under clonal growth, a widespread life history in nature, remain poorly understood. Here we show that branching events in a seagrass clone or genet leads to population bottlenecks at the cellular level and hence the evolution of genetically differentiated modules. Studying inter-module somatic genetic variation, we uncovered thousands of SNPs that segregated among modules. The strength of purifying selection on mosaic genetic variation was greater at the intra-module comparing with the inter-module level. Our study provides evidence for the operation of selection at multiple levels, of cell population and modules. Somatic genetic drift leads to the emergence of genetically unique modules; hence, modules in long-lived clonal species constitute an appropriate elementary level of selection and individuality.

---

All multicellular species, from plants to humans, are genetic mosaics, owing to mitotic errors (somatic mutations) during growth and development^1,2^. In unitary species, the resulting intra-organismal genetic heterogeneity may lead to genomic conflict and is often associated with degenerative disease such as cancer^3^ (but see ref ^4^). Somatic genetic variation may play a different, more positive role in species undergoing asexual (or clonal) reproduction, hereafter called modular species, featured by 65% of all plant families^5^ and 35% of all animal phyla^6^. Modular species have a simple body plan, and often indeterminate growth, during which iterative units (modules) emerge by asexual proliferation through fission, budding or branching^7^. Modules originating from the same zygote collectively form the clone or genet^8^. Under modular organization, somatic genetic variation may segregate when new modules are formed. Inevitably, a subset of the original proliferating cell population are precursors for any new module, and this genetic bottleneck will affect the frequency of alleles in asexual offsrping. This process of *somatic genetic drift* at the level of proliferating cell populations is fundamental and inevitable and needs to be understood before addressing selection^9-11^. Importantly, somatic genetic variation may become fixed throughout the entire module’s cell population, while other somatic genetic variation may continue as genetic mosaics (Fig. 1, Supplementary Fig. 1).

**Fig. 1.**
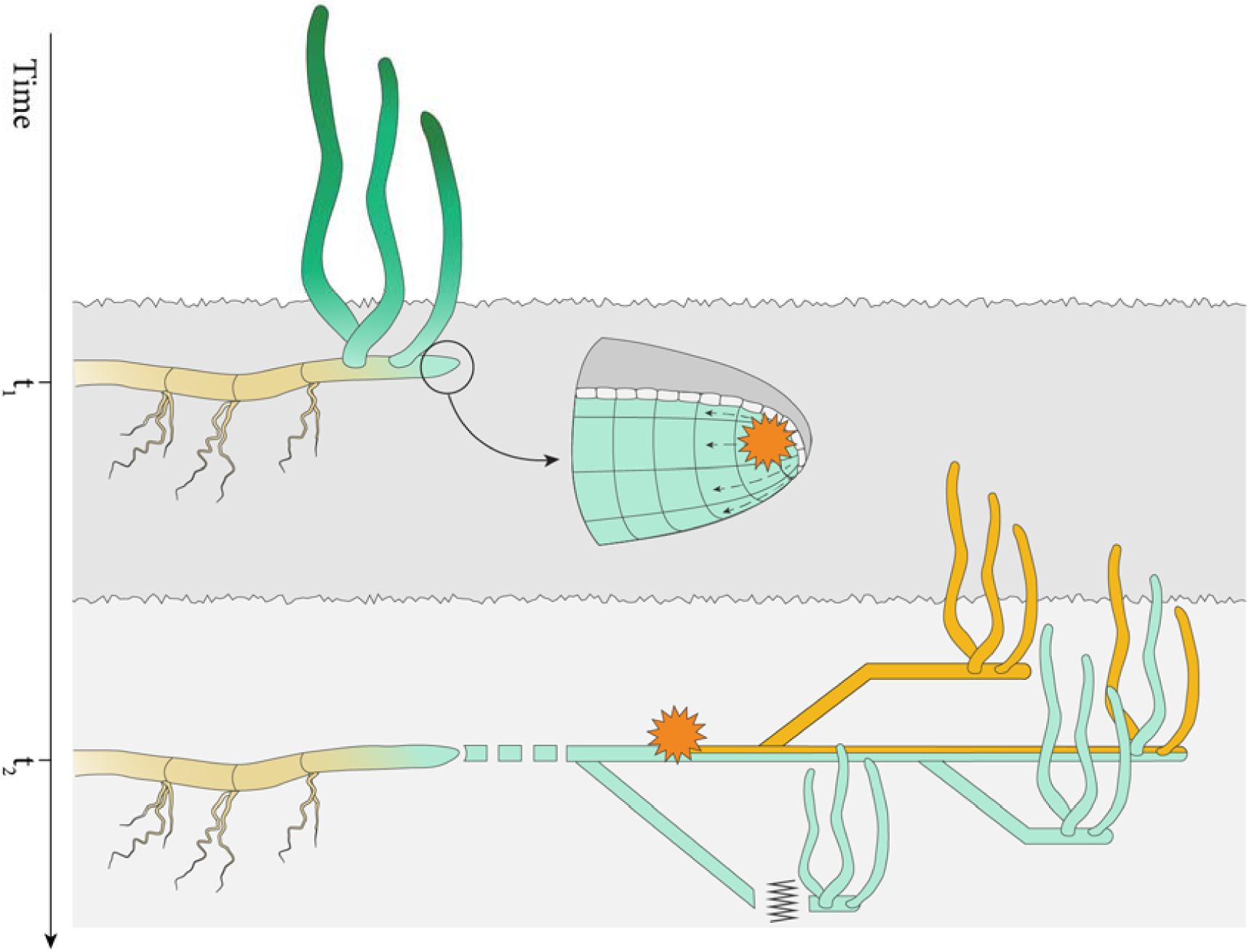
Somatic genetic drift among modules of a clone (=genet) of the seagrass *Zostera marina*. During each branching event, a finite subsample of proliferating cells forms each new module. At a given locus, a new module may remain in the mosaic state or become fixed for the novel mutation (orange), or lose the new somatic genetic variation. In *Z. marina*, modules become physically independent after a few years.

Recurrent events of somatic genetic drift thus create a third intermediate layer of genetic variation in between the cell population level (as genetic mosaicism) and sexually recombining genotypes (populations of genets), namely at the module level^6,12,13^. In line with this notion, population genetic models have demonstrated that with increasing longevity, there may be far more mitotic cell divisions during one zygote cycle than meiotic ones^14^, potentially providing large mutational input for both, selection and somatic genetic drift to operate^15,16^. Accordingly, several studies have demonstrated the emergence of phenotypic differences among modules of the same genet^10,17,18^. However, the corresponding genome-level assessments of frequency, dynamics and possible functional consequences^19,20^ of somatic genetic variation are currently lacking for any modular species.

Here we examined a species with indeterminate asexual proliferation, the seagrass *Zostera marina*, belonging to a group of marine angiosperms that is well known for their massive clonal proliferation and longevity^21-23^. The relatively compact genome of *Z. marina* has recently been characterized^24^, allowing us to re-sequence modules of a large genet with high coverage. The objective of the present study was to obtain whole-genome-level quantification of inter- and intra-module genetic heterogeneity, in order to assess the potential for intra- and inter-module selection. We tested the hypotheses that (i) the branching events (where somatic genetic drift occurs) contribute to segregating intra-module mosaic diversity into non-mosaic, fully heterozygous genotypes among modules; and (ii) that molecular evolution at the intra- and inter-modular levels differs and would provide evidence for multi-level selection.

## Abundant somatic genetic variation in a single plant genet

In order to describe genomic patterns of a putative large seagrass genet in space, twenty-four modules (ramets, ref ^8^) were collected by SCUBA in 2016 along predetermined distances along two transects at the site Angsö in SW Finland, Baltic Sea (Fig. 2a and Supplementary Table 1). This is one of the many sites that feature mega-clones of hundreds of meters in spatial extension^21^. The selected meadow (covering an area of ca. 6 ha) has been previously characterized as “clonal” based on microsatellite genotyping^21^. We estimate that at least 5,300 branching events preceded each of the contemporary living leaf shoot or modules (see Methods). Bulk DNA from basal leaf shoot tissue encompassing the meristematic region and the base of the leaves was sequenced with an average coverage of 81x per module (Supplementary Tables 2 & 3). Sequenced reads were mapped against the *Z. marina* reference genome^24^ with an average mapping rate of 94%, noting that the reference genome was from a clonal meadow only 22 km from the samples used in the present study (Supplementary Fig. 2). We focused first on fixed genetic variants that were present in all the cells within the module. A total of 38,831 fully heterozygous SNPs were detected after stringent filtering (Supplementary Fig. 2) that had a different allele compared to the reference genome.

**Fig. 2.**
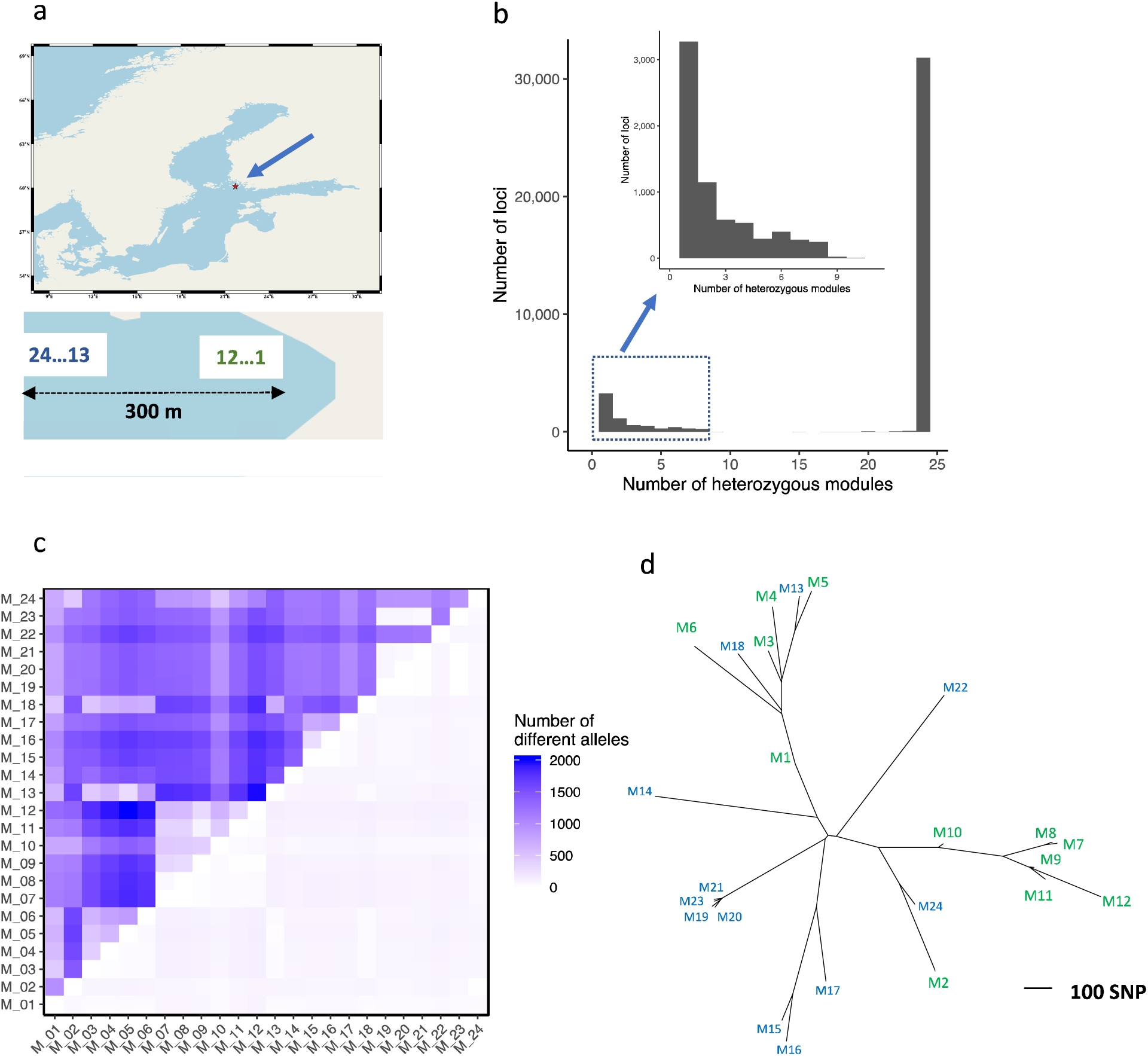
Genetic heterogeneity among the 24 seagrass modules belonging to a single genet (=clone). **a**, Location of the sampled seagrass population in southern Finland and sampling points along a transect with sample numbers running from shallow to deep **b**, Histogram of shared SNP polymorphisms among 24 modules. The x-axis depicts the number of heterozygous modules, the y-axis represents the number of shared SNPs among all samples. **c**, Pairwise number of heterozygous SNPs differing among modules. Above the diagonal: all SNPs; below the diagonal: only non-synonymous SNPs. **d**, Neighbor-joining tree based on 24 sampled modules (green are shallow, blue are deep). The tree is based on 3,095 parsimony informative SNPs (total 7,054) and calculated using 1,000 bootstrap replicates. Most nodes had a bootstrap support >95%, see Supplementary Figure 6.

The resulting multi-locus SNP genotypes were first used to assess whether or not all 24 sampled modules were members of the same genet, i.e. originated exclusively by vegetative propagation. Members of the same genet will harbor an excess of shared heterozygous loci because under mitosis, all SNPs of the ancestral zygote will remain heterozygous in offspring. The majority of loci (31,777; 81.83%) were consistently heterozygous in all 24 modules (Fig. 2b, right hand bar), confirming that all sampled modules belong to the same genet (Supplementary Fig. 3). The analysis also revealed a substantial number of SNPs that were heterozygous in a subset of modules (Fig. 2b, left-hand side). These SNPs are best explained by somatic mutations that emerged initially as mosaics but later increased in frequency to become fully heterozygous genotypes via somatic genetic drift; these SNPs were the focus of our study.

## Inter-module genetic diversity within the large genet suggest widespread somatic genetic drift

After subtracting loci that were shared among all 24 modules (blue SNPs in Supplementary Figure 1), we identified 7,054 SNPs that originated by somatic mutations (Fig. 2b). Experimental validation of a subset of the inferred SNPs by genotyping all combinations of 14 SNP loci tested in 24 modules revealed that all genotypes were accurately annotated (Supplementary Figure 4 & 5, Supplementary Table 4). Based on the SNP polymorphism observed, pairwise genetic distances were calculated and revealed an average pairwise SNP distance of 1,216 among different modules (Fig. 2c and Supplementary Table 5). Based on the genome annotation^24^ we identified 1,672 SNPs located in genic regions, and 597 in coding regions, of which 432 were non-synonymous SNPs (Supplementary Dataset 1). We then examined how the genetic similarities would correspond to the spatial location of modules. A neighbor-joining tree of the sampled modules revealed geographically close modules clustered together (Fig. 2d and Supplementary Fig. 6), confirming the mutation accumulation hypothesis predicted for large vegetative genets. There were, however, some notable exceptions (modules M02, M13, and M18). The disagreement between the branching topology and geographical location of these modules indicates that they were recently introduced, most likely by uprooting and drifting to the current location as shown for another seagrass species^22^.

Our analysis further revealed 1,654 insertion-deletion polymorphisms (indels) ranging from 1-201 bp in size (median 4 bp, mean 8.53 bp). The frequency of indels in our data is in line with other assessments of the expected ratio between substitutional mutations leading to SNPs and insertion / deletion polymorphisms^25^. The distribution of indel-based genetic diversity revealed largely similar patterns to that of SNPs with respect to the pairwise inter-module divergence and the resulting topology (Supplementary Fig. 7 & 8). This is consistent with a view that many of the somatic polymorphisms are behaving neutrally, regardless of whether these are of the indel or point mutation type. We also identified one larger structural polymorphism of 200 bp in size, and 7 microsatellite polymorphisms, which occurred along the branching history reflected in the NJ-tree topology. All observed somatic variants independently confirmed the module genealogy (Supplementary Fig. 9 & 10).

## Intra-module somatic mutations and genetic mosaicism

Any somatic mutational input enters the module at a low frequency through a single, proliferating cell and stays as genetic mosaic unless it is lost or rising to fixation (Fig. 1). To quantify somatic genetic polymorphism at intra-module allele frequencies <<0.5, we re-sequenced three modules (M08, M10, and M12) to a very high coverage of 1370x (Supplementary Fig. 2, Supplementary Table 3 & 6). An independent, restriction enzyme based method (see Methods) verified 13/15 tested mosaic polymorphisms (Supplementary Fig. 11, Supplementary Table 7 & 8). Both non-confirmed SNPs were at the lower range of the variant read frequency (*f*=0.05 and 0.06, respectively). This ultra-high coverage dataset was first used as a standard to obtain an alternative estimate of the proportion of true, fully heterozygous genotypes as described in the previous section (see Methods). Reassuringly, 81.09% (78,087/96,291) were confirmed as being fully heterozygous, demonstrating that our genotyping pipeline was largely successful in detecting fixed SNPs among modules of a single genet and distinguishing them from the mosaic ones.

We found 4,973 intra-module somatic SNPs per module on average (Fig. 3a). Of these, 71.98% were in a mosaic state. Among those SNPs, 820 were found in genic regions, and 301 in coding regions, of which 198 were non-synonymous. None of non-synonymous ones overlapped with the non-synonymous, fixed SNPs detected among modules. The distribution of variant read frequency (VRF), our proxy for intra-module allele frequency (AF), showed concordant patterns with multiple peaks within the low-frequency regions for each module, indicating the coexistence of different proliferating cell populations (arrows; Fig. 3b-d). All apical shoot meristems examined thus far in flowering plants are organized into different cell populations or layers^26,27^. Hence, the modes at *f*<0.5 likely correspond to somatic genetic variants that have come to sub-fixation in separate meristematic layers^26-28^. We also infer from the VRF distribution that cells within one meristematic layer date back to a few initial cells^29^, which explains the rapid sub-fixation of somatic mutations. A second non-exclusive explanation for the pronounced modes of the VRF distribution would be intra-module positive selection, which may favor a particular cell lineage possessing a driver mutation with which an entire cohort of neutral background mutations would “hitchhike”^30^.

**Fig. 3.**
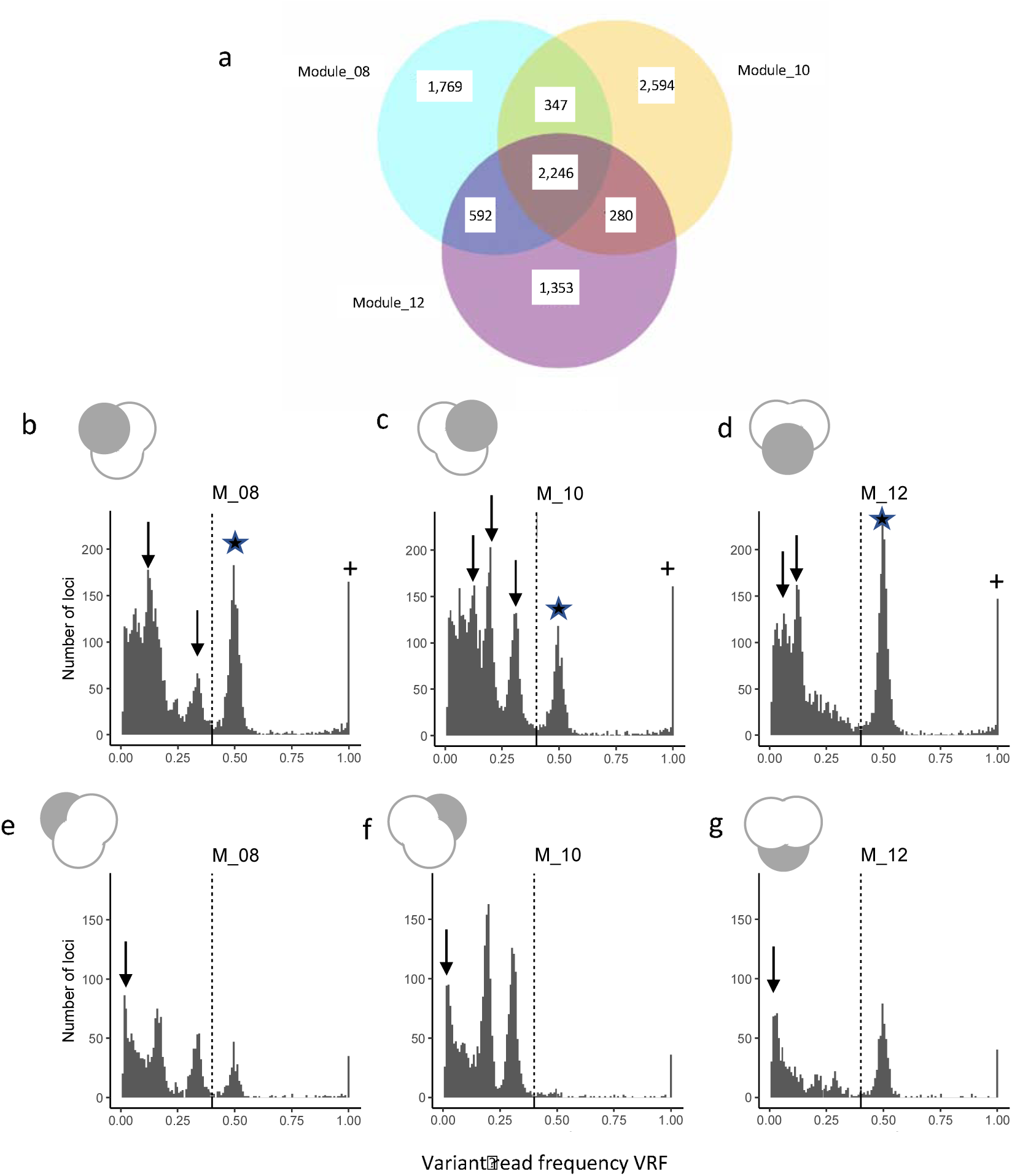
Intra-module somatic genetic variation. **a**, Venn diagram depicting overlap of mosaic and non-mosaic SNP variants in three ultra-deep sequenced modules (1370x) that emerged via somatic mutation. In all panels, a miniature Venn diagram indicates which sample of SNPs was considered. **b-d**, Histogram of intra-module variant read frequency (VRF) for Module_08, Module_10, and Module_12. The x-axis represents the variant read frequency as a proxy for the variant allele frequency, and the y-axis represents the number of SNPs. SNPs were categorized into three types: (i) non-fixed, intra-module mosaic-type SNPs in which arrows depict differentiated cell populations (ii) fixed heterozygotes (peak at *f*=0.5, star); and fixed homozygotes (“+”, somatic mutation 0/1- >1/1; peak at *f*=1). **e-g**, VRF histograms for the private variants in each module. Here, arrows indicate the mutational input, following a power law accumulation. In all panels, the dashed line depicts the threshold of VRF below which a SNP was considered to be in a mosaic, non-fixed state.

Under exponential clonal expansion of modules, the accumulation of neutral genetic variants should follow a power-law distribution, leading to a left-hand peak of rare variants^31,32^. Assuming that many of the intermediate modes of the VRF distribution represent somatic genetic variants that were fixed within confined meristematic layers, we excluded those SNPs shared among modules. When plotting the distribution of module-specific somatic genetic variants only, the expected left-hand peak of variants appeared, indicative of neutral mutational accumulation proportional to 1/*f* as described in modeling studies (Fig. 3e-g).

## Effect of purifying selection at the intra-module versus inter-module level

Given the predominance of homozygous sites in the ancestral zygote (99.91%, see Methods), most somatic mutations will result in a homozygous-to-heterozygous transition. The fitness effects of any mutation depend upon their degree of dominance and tissue-specific expression^19^. While many deleterious mutations will be fully recessive and only expressed in haploid gametes^33^, a fraction has been shown to be partially dominant and can be subject to selection even when heterozygous^34^. Hence, we compared the strength of purifying selection at the intra-module (i.e., cell population) and inter-module levels. The estimation of intra-module selection was based on mosaic SNPs identified in three modules (1370x data; module M08, M10, and M12). Assessments of inter-module selection were based on fully heterozygous SNPs among all 24 modules (∼80x data). We found significant signals of purifying selection at both inter- and intra-module level (Fig. 4). The dN/dS ratio indicated weak purifying selection at the inter-module level, while this signal was much enhanced at the intra-module level (both for module-specific mosaic SNPs and all mosaic SNPs) (Fig. 4a). The frequency of somatic genetic variation was considerably depleted in genic loci (Fig. 4b) and within coding sequences (Fig. 4c) relative to the entire genome. Furthermore, there was no significant difference comparing the frequency of SNPs in coding sequences (CDS) relative to all genic locations at both selection levels (Fig. 4d), indicating that several genic mutations may also be subject to purifying selection when located in untranslated regions or introns. In none of the variables tested did we find a difference among all mosaic SNPs vs. those that were module-specific.

**Fig. 4.**
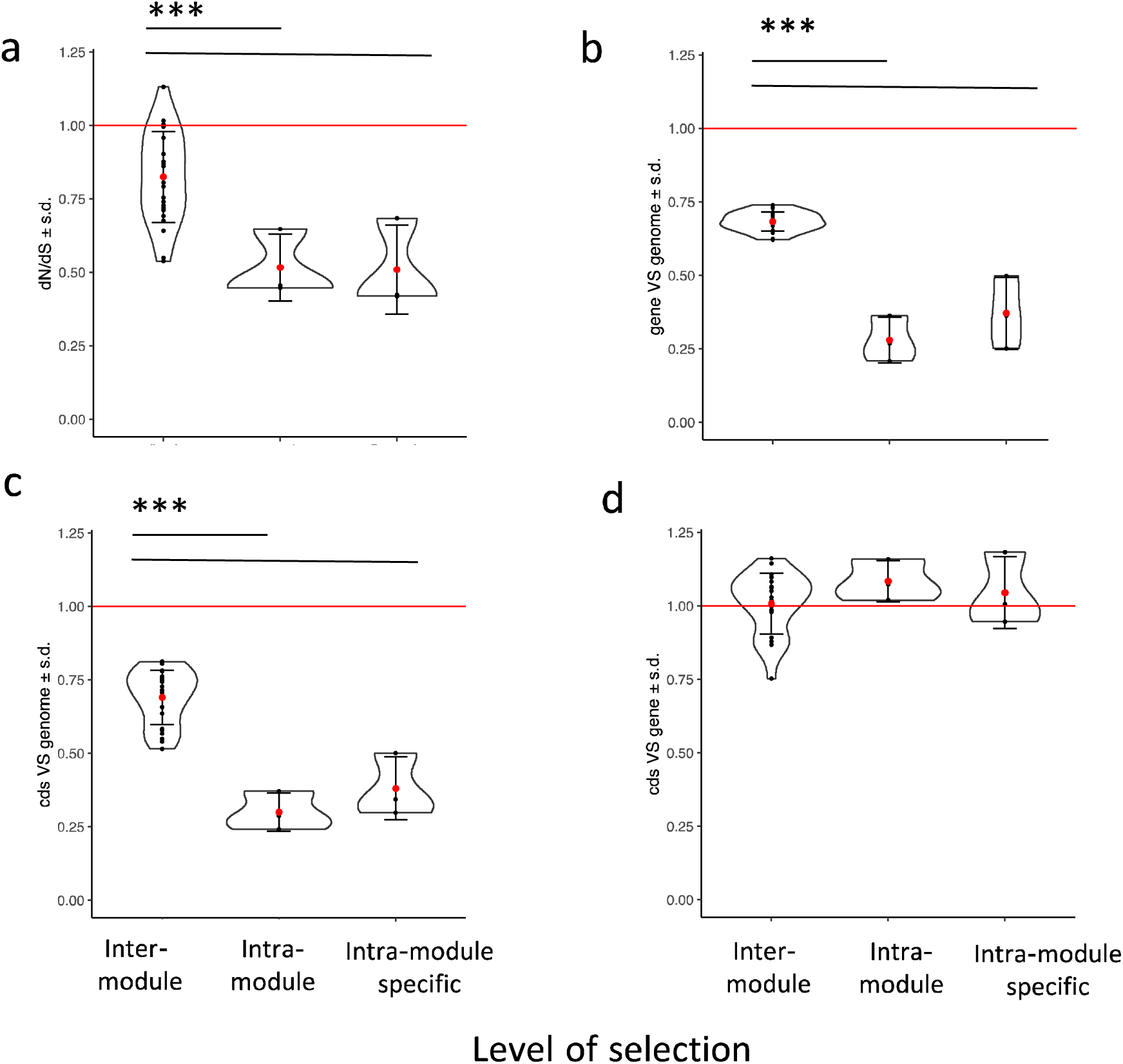
Comparison of purifying selection at the inter- and intra-module level. In each panel, three levels of selection are distinguished, (i) among modules based on fully heterozygous, fixed SNPs; (ii) within modules M08, M10 and M12 (i.e., in mosaic status) based on either all SNPs or only those SNPs that are not shared; and (iii) specific to a single module (cf. Venn Diagram, Fig. 3a). In all panels, means ±1SD are given; the red line represents the expected value based on selective neutrality. Significant differences among the levels of selection are indicated with asterisks (***p<0.001). **a**, dN/dS calculated based on intra-module somatic mutations. All three values significantly deviated from neutrality (Supplementary Table 9). **b**, Fraction of genic SNPs compared to the entire genome, standardized for the abundance of genic regions. **c**, fraction of SNPs in coding regions compared to the entire genome, standardized for the proportion of coding regions. **d**, Fraction of coding SNPs within genic regions (intron+exon+UTR).

Our results indicate that selection in clonal seagrass operates at multiple levels. First, we found that purifying selection was stronger among somatic genetic variation still at the low-frequency mosaic state, i.e., when it occurs among cells, while relaxed among modules (Fig. 4a-c). This was predicted in earlier models suggesting that selection at the cell level within the module is effective in purging the asexual population from deleterious mutations^11,35-37^, especially when the meristem has a layered structure^27^. Additional evidence for multi-level selection came from the comparison of gene function. The fully heterozygous SNPs compared to intra-module, mosaic SNPs were each associated with different non-overlapping functions. The former polymorphisms were located within genes enriched (among others) for the molecular function “protein binding”, the cellular component “nucleus” and the biological process “cellular protein modification”. The latter mosaic SNPs were situated in genes significantly enriched for the non-overlapping GO terms “structure of cytoskeleton” (molecular function), “protein microtubule” (cellular component), “protein phosphorylation” (biological process) (all *P*-values <0.005; Supplementary Table 9).

## Dynamics of intra-module allele frequency among modules

The intra-module somatic mutations were detected independently for each of the three modules. Of a total of 9,208 unique loci, 62% (5,743) were specific to one module, followed by 24% (2,246) that were shared by all three modules (Fig. 3a), and 14% shared by pairs of modules. The latter two data categories provided an additional independent verification that our SNP calling within modules was accurate, as the detection of somatic genetic variants was technically independent among the three modules. We used the inferred genealogy to assess the dynamics of intra-module allele frequency among modules M08, M10 and M12 (Fig. 2d). Under asexual propagation, the somatic mutations accumulate along the growth path of the rhizome. Accordingly, the more recently two branches have diverged, the more similar intra-module allele frequencies should be at a given locus. With a deep coverage of ∼1370x, variant read frequency (VRF) should be a reasonable proxy for corresponding allele frequencies (AF), following a binomial distribution determined by both intra-module allele frequency and read coverage at the given locus^38^. We therefore set a conservative coverage threshold of 500x as prerequisite to estimate allele frequencies via VRFs. We focused on the 2,246 somatic genetic variants shared by all 3 modules, and visualized their normalized frequencies in a ternary plot (Fig. 5a). Most of the loci were found in the center, indicating similar VRFs among all three modules, a finding which is in accordance with long-term stability of periclinal genetic mosaics (or chimeras) in horticultural plants^39^. Apparently stable frequencies also applied to those shared by a pair of modules (Supplementary Fig. 12a-c).

**Fig. 5.**
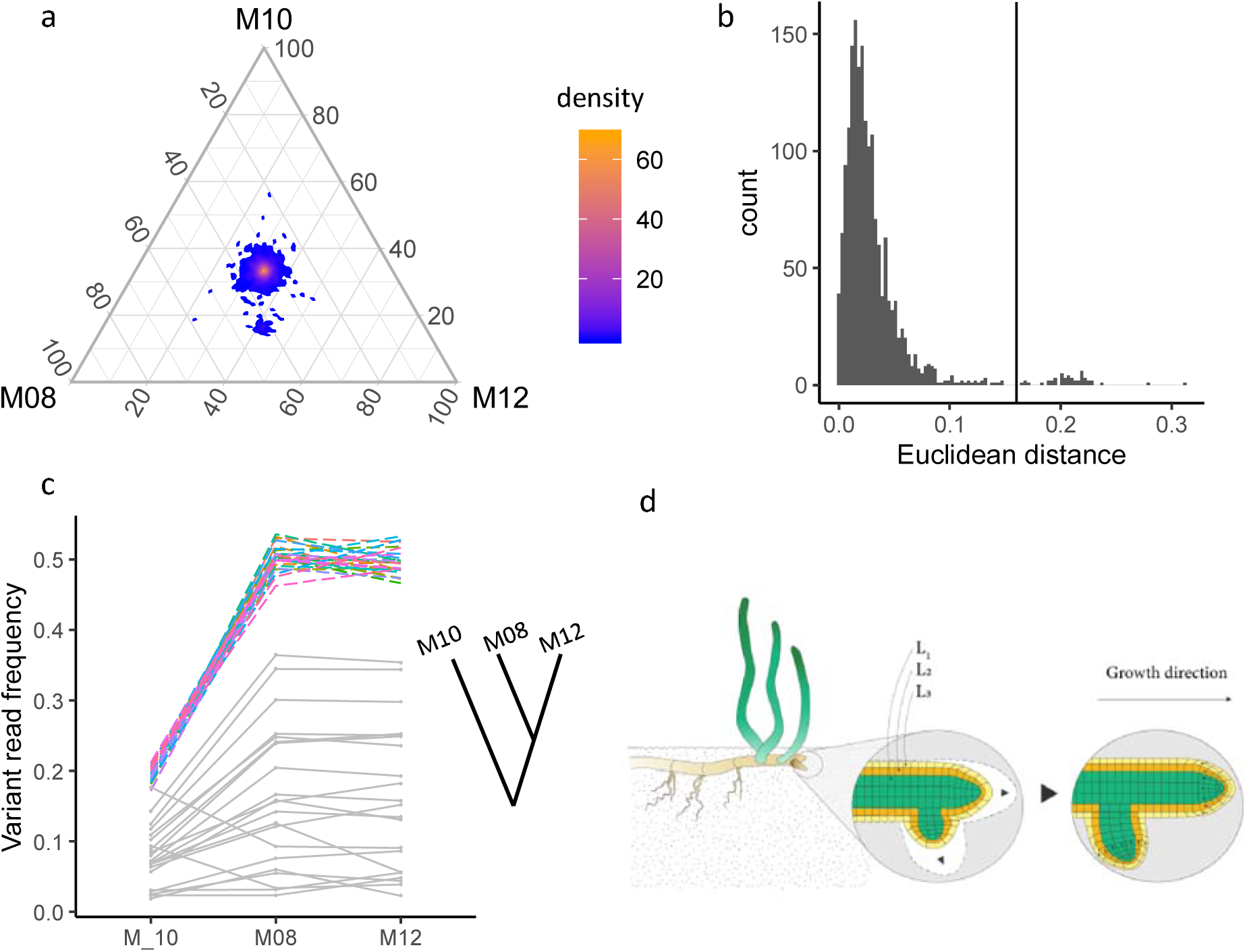
Dynamics of intra-module somatic mutations among the three modules. **a**, Ternary plot comparing intra-module variant read frequencies present in all three modules (cf. center in Venn diagram Fig. 3a). **b**, Histogram of differences in variant frequency of SNPs to the center of the ternary plot. The vertical line depicts the threshold above which loci are considered to have significantly changed in frequency. **c**, using this threshold, a total of 47 SNPs changed in frequency, in line with the reconstruction of branching events (see inset: M10 is at the genealogical base of the branching of M08 and M12, cf. Fig. 2d). **c**, Variant read frequency (VRF) across the three modules (M10, M08 and M12) based on 47 SNPs meeting the threshold in panel b. The 27 SNPs that jointly came to fixation are shown in color. **d**, diagram illustrating how the frequency of intra-module mosaics is faithfully transmitted across the hypothetical cell population layers L1… L3 during branching.

In the ternary plot, there were also some clear outliers, indicating different allele frequencies among some shared SNPs, on which we focused next. When examining the histogram of individual SNP distances to the center (Fig. 5b), a discontinuity at ∼0.16 was set as threshold beyond which loci were considered to have changed in allele frequency. Of the resulting 47 SNPs (Fig. 5c) most showed a consistent pattern in concordance with the topology revealed by the NJ tree (Fig. 2d). Module M08 and M12 recently diverged; accordingly, they revealed similar frequency dynamics that jointly differed from those of module M10 (Fig. 5c). More importantly, 26/47 of VRFs in module M08 and M12 went from a mosaic to the fully heterozygous state (i.e. *f*≈0.5), coming from a lower frequency in the ancestral M10 where the same SNPs were still in a low-frequency state. Since ancestral (M10) and derived frequencies (M08 and M12) of these 26 loci behave largely similarly, the most parsimonious explanation is that they became fixed in one single asexual sweep driven by drift, selection or both. Accordingly, our results demonstrate the rise in somatic genetic variation to a fully heterozygous state among all cells of a module. While cell populations in meristematic layers in angiosperms are separated^26-28^, sometimes cells can change among layers, otherwise it would be impossible for low frequency somatic variants emerging in one layer to come to a full heterozygous state^39^. Results of 347, 592, and 280 SNPs shared among pairs of modules (M08 – M10, M08 – M12 and M10 – M12), respectively, show a similar pattern of most VRFs to stay constant, while only a small fraction changed in allele frequency (Supplementary Fig. 12a-c).

## Discussion

A modular organization including asexual budding, branching or fission is a widespread life-history trait shared by plants, animals and fungi^5,6,40^. In the present study, we trace how somatically generated variation that is initially in a mosaic state segregates into differentiated module genotypes within a large genet of a seagrass (Supplementary Figure 13), a group of marine plants that is known for its very large clonal expansion. The amount of somatic genetic variation detected among modules is roughly 100 times higher than that found among branches of long-lived trees such as oak^41,42^ (Supplementary Table 10). We argue that this is caused by the high number of branching events that are not restricted by maximal tree size. Modular species with indeterminate clonal growth such as corals, reed, ferns, aspen, fungi or seagrasses, in contrast, can grow, branch, and fragment as long as space for expansion is available. Each branching, in turn, presents an opportunity for somatic genetic drift, eventually resulting in strong genetic heterogeneity among different modules. It has been speculated as to whether and how long-term vegetative growth may lead to senescence in clonal plants^43,44^. In the northern Baltic Sea, *Z. marina* usually does not complete its sexual life cycle, but it is unclear whether this is driven by too short a growing season or genetic factors. Notwithstanding, given that a conservative age estimate for the genet studied here ranges from 750 - 1,500 yrs (with an upper bound of 6,000 yrs) our results are in agreement with other studies that only found a significant decline in male fertility of aspen genets >4,000 yrs old^44^.

Our findings confirm the predicted presence of multiple levels of selection in modular species^16,28,36^. This was evident through different strength of purifying selection acting on mosaic polymorphism within modules versus somatic genetic differentiation among modules. Considering that 432 of the fixed SNPs were nonsynonymous, our study suggests the potential of somatic genetic variation as a contributor to molecular adaptation (Supplementary Dataset 1). Detecting asexual selective sweeps is difficult as any favorable mutation will be dragged along with its entire mutational background behaving as one large genetic linkage group in the absence of recombination^30,32^. A critical next step will be to test for positive selection^32^ at the molecular level and to study possible adaptive consequences of somatic genetic variation in genets of *Z. marina*.

Once haplotypically fixed, somatic genetic variation will also enter the sexual cycle in flowering plants^2,19^, while this is debated in basal metazoan species^45^. Notwithstanding the sexual cycle, our data support the view that under asexuality, modules are the appropriate elementary level of selection and individuality as the rapid accumulation of genetic variation through somatic genetic drift makes them genetically unique^46^. Whether or not the identified inter-module genetic differentiation translates into different phenotypes is an open and highly intriguing question^4,12,13,19,47^. Experimental studies have revealed differences in physiological performance among asexually propagating modules of clonal plants and algal thalli^10,17,18,20^, suggesting the principal significance of inter-module phenotypic selection. We suggest that somatic genetic drift producing inter-module genetic differentiation will apply to many clonally proliferating species, providing an additional source of genetic variation for adaptation. This includes, e.g. fungi, clonal trees such as aspen and basal animals such as reef-building corals, hydrozoans, bryozoans, colonial ascidians and any other invertebrate species with indeterminate growth and module formation^6,40,47^. A comparative view across the animal, plant and fungal kingdom as to the abundance, fate and selective consequences of somatic genetic variation is thus highly warranted.

## Supporting information

Supplementary_Figs_Tables_all

## Acknowledgements

This study was supported by a 4-year PhD scholarship from the China Scholarship Council to L.Y. We thank Jeanine L. Olsen and Benjamin Werner for helpful comments on earlier versions of the manuscript. Thanks to Susanne Landis for creating some of the illustrations. Sampling permit # MH 5448/2015 was granted through Parks and Wildlife Finland (Metsähallitus).

## Methods

### Study species, site and sampling design

Our study species belongs to the seagrasses, a polyphyletic group of angiosperms (flowering plants) that secondarily returned to the sea^24^. *Zostera marina* L. (eelgrass) is the most widespread marine angiosperm of the northern hemisphere, occurring from subtropical areas to subarctic areas in the Atlantic and Pacific Ocean. The studied eelgrass *Z. marina* bed is located off Ängsö Island in the Archipelago Sea, southern Finland (Fig. 2a), covering an area of 300 m × 200 m (=6 ha). As in all seagrasses, in *Z. marina*, physiologically independent modules (or “ramets” sensu ref ^8^) emerge naturally since rhizome connections are disintegrating in nature after a few years (Fig. 1). Initial microsatellite screening with 8 markers revealed 3 multi-locus genotypes that were only distinguishable at one locus each^21^. However, using whole genome re-sequencing in this study we found only one genet (Fig. 2b), indicating that the results based on limited number of microsatellite markers can be biased by high level of somatic mutations (Supplementary Figure 10). Taking *Z. marina* module density of 311 modules m^−2^ (ref^48^), a conservative estimate of areal extent of 90,000 m^2^ yields 27,990,000 modules. Assuming branching with constant bifurcation and no mortality, thousands of branching events (27,990,000^0.5^ ≈ 5,300) preceded each of the existing modules. As for the age estimation, assuming 3 branching events per year, we arrive at 5,300/3 ≈1,800 yrs. The spatial extent allows for an alternative estimate based on linear dimension. Assuming a horizontal spread of about 20 cm yr^−1^ (ref ^49^) and no intermittent mortality, an expansion to 300 m can be reached in between 750-1,500 years, assuming the ancestral module in the center /peripheral to the meadow, respectively. For an absolute upper age limit we assume 6,000 yrs, the time when the current salinity levels were reached after the re-connection of the then Baltic freshwater lake to the ocean^50^. Twenty-four modules were collected by SCUBA in 2016 along transect with a shallow and a deep section (Fig. 2a, Supplementary Table 1). Twelve modules each were collected along predefined distances at approximately 2 m and 4 m water depth, respectively.

### Whole-genome resequencing

Bulk DNA of the meristematic region and the basal portions of the leaves, weighing approximately 20-50 mg (fresh mass) was extracted using NucleoSpin Plant II kit (Macherey-Nagel, Germany). DNA concentrations were determined using a Qubit Fluorometer (Thermo Fisher Scientific) and Nanodrop Spectrophotometer (Thermo Fisher Scientific). We checked the integrity of the genomic DNA by agarose gel electrophoresis against a molecular weight (mw) standard and always detected a crisp band at >20 kb in mw. DNA was sent to IKMB (Institute of Clinical Molecular Biology, University of Kiel, Kiel, Germany) on dry ice for library construction and sequencing preparation. TruSeq Nano DNA libraries were constructed for each of the 24 samples with insert size ranging from 479 bp to 515 bp. All the 24 libraries were sequenced on Illumina HiSeq4000 platform at a targeted coverage of ∼80x. Based on the topology of a NJ tree, three modules were selected (Modules 08, 10, and 12) to be ultra-deeply re-sequenced on Illumina NovaSeq6000 platform at a targeted coverage of ∼1000x.

### Sequence processing and filtering

The quality of Hiseq paired-end reads (2 × 151 bp) and Novaseq paired-end reads (2 x 151 bp) was assessed by FastQC v0.11.7 (Ref^51^). Reads were then filtered and trimmed using Trimmomatic v0.36 (Ref^52^). Adaptor contaminations were trimmed, and reads with leading or trailing Phred score <20 were removed. The reads were also filtered with a sliding window of size 3; the threshold of average Phred score was set to 15. Trimmed reads with length <36 bp were also removed. FastQC was used for a second round of quality evaluation on the clean reads. After having been trimmed and filtered, the clean Hiseq reads had the average coverage of 81x per sample, and the clean Novaseq reads had the average coverage of 1370x per sample. Clean reads were then mapped against the *Z. marina* reference genome v2.1 (ref^24^) using BWA-MEM v0.7.17 (Ref ^53^) with default parameters. The aligned reads were sorted using SAMtools v1.7 (Ref ^54^), and duplicated reads were marked using MarkDuplicates tool in GATK v4.0.1.2 (Ref ^55^).

### Detection of fully heterozygous, fixed SNPs

*Z. marina* is a diploid plant with no evidence for recent genome duplication^24^. Given the predominance of homozygous sites in the ancestral zygote (proportion of heterozygous loci in the ancestral zygote, 0.09%, see calculation below), most somatic mutations will change a homozygous to a heterozygous state, notwithstanding the rare case where a heterozygous locus will change to a homozygous state (Supplementary Figure 1). Another case would be even rarer where the same mutation double hits the same homozygous locus and change it to a different homozygote. Thus, within a given cell, we assumed the variant allele to be either present once or none. Once a variant allele is present in all the cells of a module, we considered the somatic polymorphism to be fully heterozygous, or “haplotypically” fixed (Fig. 1a, Supplementary Figure 1), which will in an expected intra-module allele frequency of exactly 0.5. SNPs were called by GATK v4.0.1.2 package. HaplotypeCaller was used to generate general variant calling files (gvcf), which were then combined by CombineGVCFs into a single gvcf file. The merged gvcf file included 81x coverage data for all 24 modules and 1370x data for modules M08, M10, and M12. GenotypeGVCFs was used to run joint genotyping.

We first applied GATK hard filtering to filter the SNPs (Supplementary Figure 2). The density plots of the recommended annotations were drawn with ggplot2 (Ref ^56^), based on which the thresholds were decided: QualByDepth (QD < 15.0), FisherStrand (FS >10.0), RMSMappingQuality (MQ < 40.0), MappingQualityRankSumTest (MQRankSum < −1.5), and DP (DP > 8000.0). DP > 8000.0 was aimed to remove the SNPs potentially caused by genomic duplication, which was featured with higher coverage. We then used VCFtools v0.1.15 (Ref ^57^) to remove genotypes with GQ < 30 (--minGQ 30) or DP < 20 (--minDP 20). After having removed these genotypes, VCFtools was used to remove SNPs with more than 2 alleles (--min-alleles 2 --max-alleles 2), SNPs with minor allele frequency < 0.01 (--maf 0.01), and SNPs with more than 4 missing genotypes (--max-missing-count 4).

### Detection of fixed indels

Indels were also called by GATK v4.0.1.2 package. We first applied GATK hard filtering to filter the indels: QualByDepth (QD < 15.0), FisherStrand (FS >10.0), and DP (DP > 8000.0). We then used VCFtools v0.1.15 to remove genotypes with GQ < 30 (--minGQ 30) or DP < 20 (--minDP 20). After having removed these genotypes, VCFtools was used to remove indels with more than 2 alleles (--min-alleles 2 --max-alleles 2), indels with minor allele frequency < 0.01 (--maf 0.01), and indels with more than 4 missing genotypes (--max-missing-count 4).

### Verification of fully heterozygous (=haplotypically fixed) SNPs

We developed an independent, non-sequencing based SNP verification method using restriction enzymes with motifs specific to the polymorphic site (Supplementary Fig 2). Accordingly, we searched the reference genome for RE motifs encompassing SNPs. Upon designing fluorescence-labelled primer pairs, a ∼400 bp PCR fragment surrounding the target SNP was amplified, which was subsequently cut by restriction enzymes. Fragments carrying identical sequences to the reference genome possessed intact restriction sites and were cut, while fragments with a variant allele lost the restriction sites and remained uncut. Fluorescently labelled PCR amplicons were run on Applied Biosystems 3100xl sequencer using the fragment analysis module. The heterozygote would have fluorescence peaks at two different locations. Reassuringly all tested SNPs were verified, i.e. fourteen SNPs in combination with all 24 modules (Supplementary Figure 5). In some instances, we could not obtain clean PCR products using the designed primer pairs which prevented the subsequent SNP verification.

We then verified if inter-module SNPs were truly fully heterozygous (i.e. not occurring in some lower, mosaic frequencies), using the three 1370x sequenced modules M08, 10 and 12. Under fixation, the intra-module allele frequency of both the variant and the ancestral allele is *f*= 0.5. With a coverage of 1370x, the read frequency calculated based on the Novaseq dataset was taken as proxy for intra-module allele frequency. Our analysis was restricted to those heterozygous genotypes where the coverage was >500x. Confidence intervals were calculated according to a binomial distribution of the variant read frequency centered at 0.5, and with a standard deviation (SD) determined by the coverage (Ref ^58^). As an example, when coverage=500, SD would be SD=0.022, two times which was set as the threshold to obtain the ±95% confidence interval. Accordingly, any heterozygous genotype with a VRF in the confidence range [0.5±2*0.022] was considered fixed.

### Verification of modules belonging to a single genet

Based on the high-quality SNPs, we plotted a histogram to assess whether or not the modules belong to the same genet, i.e. originated from a single zygote via vegetative propagation. Under mitosis the heterozygous loci in the ancestral zygote will remain heterozygous in all offspring cells and modules, and become visible as one predominant mode of heterozygous loci shared by all 24 modules. We plotted the number of loci against the number of modules sharing a particular heterozygous locus. The few missing genotypes (four missing genotypes at most, see filtering step above) were counted as being heterozygous, so that the number of heterozygous modules would be 24 as long as all the available genotypes were heterozygous. The same analyses were also done based on small insertion /deletion polymorphism (indels).

### Estimation of the number of heterozygous loci in the ancestral zygote

Under mitosis, the heterozygous loci in the ancestral zygote will be passed on to all the offspring. The raw SNP panel (before any filtering) from the GATK calling pipeline was used to estimate the number of heterozygous loci in the ancestral zygote (Supplementary Figure 2). Any SNP with missing genotypes <13 and all the available genotypes were heterozygous, were considered as heterozygous loci in the ancestral zygote.

### Inter-module genetic heterogeneity and spatial genetic structure

All SNPs with only one common genotype shared by all 24 modules were removed as these represented the genetic differences between the reference genome and the ancestral zygote which were passed on to all analyzed modules (Supplementary Fig. 1, blue). The remaining SNPs represented the somatically derived polymorphisms within the genet. The somatically derived indels were obtained in the same way. Based on the SNPs, pairwise genetic distances between each pair of modules were calculated and visualized using ggplot2 (heatmap plot). The genetic distance was quantified by the number of different alleles. SnpEff (Ref^59^) was used to annotate the SNPs. Based on the annotated nonsynonymous SNPs, the pairwise genetic distances between each pair of modules were also calculated and visualized. Based on the fully heterozygous somatic SNP panel, plot.phylo function in the ape package (Ref^60^) for R (Ref^61^) was used to construct a neighbor-joining tree to examine how the genetic similarity corresponds to spatial sampling pattern. Bootstrap support (1,000 times) was obtained using the aboot function in the poppr package (Ref^62^) for R. The same analyses were also done based on indels.

### Detection of structural variation

In order to detect larger structural variants, we applied CNVnator v0.3.3 (ref^63^) to find target loci followed by IGV v2.7.0 (ref ^64^) for a visual check. We first selected 3 samples (modules M05, M07, and M20) representing the largest distances within the NJ tree. Subsequently, CNVnator was used to call structural variations for each sample. We focused on deletions relative to the reference genome, considering that duplications were much more difficult to verify. A custom-made Python3 script was used to convert the CNVnator output to bed file format. Bedtools v2.27.1 (ref ^65^) was used to find the overlap between M05 and the other two samples. The locus was marked as target if it was called as a deletion in M05, and was missing in at least one of the other two samples. IGV was used to check if the deletion (relative to the reference genome) was true and showed polymorphism among the three modules. If both requirements were met, it was further checked among all 24 modules.

### Detection of intra-module somatic SNPs

In order accurately assess the level of intra-module somatic polymorphisms, we used the 1370x ultra-deep resequencing in modules M08, M10 and M12 to quantify mosaic genetic variation. Mosaic polymorphisms are only present in a subset of cells within the module (Supplementary Figure 1). Hence, the novel variant allele has the intra-module allele frequency lower than 0.5, depending on how many cells possessing the variant allele. As reference, we took the reference genome that was sampled ∼22 km distance from the studied location. The SNP calling was run independently for each of the three modules using Mutect2 (Ref^66^). The SNPs were filtered by FilterMutectCalls following the recommendations of the authors. Since the control sample is not equal with the ancestral zygote, the calling results included the genetic differences between the control sample and the ancestral zygote which should be shared by all 24 analyzed modules. These loci were located based on the raw SNP panel (before any filtering) from the GATK calling pipeline (Supplementary Figure 2). Any SNP with missing genotypes <13 and all the available genotypes were same, were considered as genetic differences between the ancestral zygote and the control sample, thus 215,518 loci were checked and removed, if present in the Mutect2 calling results.

### Verification of intra-module somatic SNPs

We also verified the intra-module variation using a sequencing-free, restriction enzyme based method. During the verification of fully heterozygous, fixed SNPs above we found that even the wild-type genotypes (both alleles identical to reference genome) were not digested completely, leaving a small fluorescence signal. In order to reliably detect mosaic SNPs, incomplete digestion would hence spuriously suggest low-frequency variants. Therefore, we searched for sites encompassing RE motifs where the heterozygous variant allele converted a non-restriction site to a restriction site. Thus, after digestion shorter fragments will only appear if the variant allele is present, an observation that cannot be biased by undigested wild-type alleles. We verified 15 cases in total (3 cases for M08, 4 cases for M10, and 8 cases for M12) (Supplementary Figure 11, Supplementary Table 7 & 8).

### Fixed vs. mosaic somatic polymorphisms

We estimated intra-module allele frequency, using the variant read frequency (VRF) calculated from the ultra-deep Novaseq resequencing dataset. Under 1370x coverage VRF should be a reasonable proxy for allele frequencies, and we plotted the histogram of VRF for each module. To detect the pattern of recent mutational input at low allele frequencies, we also plotted histograms only with module-specific somatic genetic variation (Venn diagram, cf. Fig. 3a). We then distinguished mosaic from fixed somatic genetic variation. The mosaic polymorphisms included both above and below fully heterozygous SNPs present among all cells of a module. For the mosaic loci, we focused on those with frequency <0.5, i.e. those below fixation in one haplotype among all module cells. According to the histograms, such fully heterozygous SNPs centered around a VRF of 0.5. There was a clear break at VRF=0.4 that separated the low-frequency SNPs, which was set as the threshold for distinguishing between low-frequency and fixed SNPs (Fig. 3c,d,e; dashed line).

### Detection of molecular selection at the intra- and inter-module level

Modular species such as *Z. marina* may be subject to two different levels of selection, among cells within modules, and among differentiated modules. We took the 7,054 fixed SNPs identified using the 81x coverage dataset to study inter-module selection. Given the predominance of homozygous sites in the ancestral zygote (99.91%), most somatic mutations cause a homozygous-to-heterozygous change. We first removed the rare cases where the locus possessed two different homozygous genotypes (0/0 + 1/1). For the remaining loci, we assumed that they originated from homozygous-to-heterozygous mutations. This data set was compared to the intra-module, mosaic SNPs identified using ultra-deep (1370x) sequencing that are representative of selection among different cell lineages within the module. SnpEff was used to identify nonsynonymous and synonymous changes. To calculate the dN/dS ratio, we also had to know the total number of nonsynonymous sites and synonymous sites in the genome, which were calculated based on the CDS sequences of the reference genome^24^. Although applying dN/dS ratio at population level has been questioned^67^, the focus of our analysis are the different intensities of selection which were significantly different at the inter- and intra-module level. The same SNP panels were also used to analyze the distribution of mutations in the genome. They were categorized into being located in CDS, gene, or other locations in the genome. We calculated (i) the ratio of mutations distributed in genes relative to the whole genome (ii) the ratio of mutations distributed in CDS relative to the whole genome (iii) the ratio of mutations distributed in CDS relative to genes, and compared the levels of selection using pairwise *t*-tests for unequal variances. Quantities were normalized based on the total length proportion of each of the genomic categories, e.g. for the ratio of mutations distributed in CDS relative to the whole genome, we calculated the ((number of SNPs in CDS)/(total number of base pairs in the CDS)) / ((number of SNPs in the whole genome)/(total number of base pairs in the whole genome)). A custom-made Python3 script was used to calculate the transition:transversion ratio between the reference allele and variant allele based on the combination of fixed polymorphisms among the 24 modules. At each level, the combination of the nonsynonymous mutations (inter-module, combination of SNPs of 24 modules; intra-module, combination of SNPs of 3 modules) served as input for the GO enrichment analysis which was performed using R 3.6.1 software and the topGO 2.36.0 package^68^. The *weight* algorithm and Fisher’s exact test were used to detect significant enriched GO terms and show GO graph topology. We followed the recommendation of topGO authors to interpret the *p*-values as corrected or not affected by multiple testing. The GO term analysis was explorative, hence the threshold for statistical significance was set at α=0.01 based on recommendations^68^. The required gene universe was obtained from the *Z. marina* genome^24^.

### Dynamics of somatic genetic variation among modules

We intended to demonstrate somatic genetic drift directly using the phylogenetic history of modules. To do so, we followed the time course of specific somatic polymorphisms, in particular whether and how often they rise to fixation. Although all three 1370x sequenced modules M08, M10 and M12 share the same zygotic ancestor, the NJ tree topology revealed module-specific paths of specific and shared somatic polymorphism. First, we identified somatic SNPs shared by all three modules, setting a minimal coverage of 500x. For each locus, we normalized its frequencies in the three modules based on the sum of the three frequencies (e.g. normalized vrf_08 = vrf_08/(vrf_08 + vrf_10 + vrf_12)). This was basis for a ternary plot where the loci with identical frequencies among the three modules are shown at the center (Fig. 5a). For each SNP, the distance to the center of the ternary plot (all normalized VRFs = 0.33) was determined. A frequency distribution of the resulting distances was plotted and a threshold was set (Fig. 5b) to identify candidate loci where allele frequencies changed, which yielded 47 loci in total. Subsequently, the VRFs of these loci across modules M08, M10 and M12 were plotted and frequency changed along the inferred genealogy was qualitatively examined (Fig. 5c). The analogous procedure was also performed for somatic polymorphisms shared by two modules (Venn diagram, cf. Fig. 3a) (Supplementary Figure 12).

## References

1 Behjati, S. et al. Genome sequencing of normal cells reveals developmental lineages and mutational processes. Nature 513, 422 (2014).

2 Wang, L. et al. The architecture of intra-organism mutation rate variation in plants. PLOS Biology 17, e3000191 (2019).

3 Frank, S. A. Somatic evolutionary genomics: Mutations during development cause highly variable genetic mosaicism with risk of cancer and neurodegeneration. Proceedings of the National Academy of Sciences 107, 1725 (2010).

4 Pineda-Krch, M. & Lehtilä, K. Costs and benefits of genetic heterogeneity within organisms. Journal of Evolutionary Biology 17, 1167–1177 (2004).

5 Honnay, O. & Bossuyt, B. Prolonged clonal growth: escape route or route to extinction? Oikos 108, 427–432 (2005).

6 Buss, L. W. Evolution, development, and the units of selection. Proceedings of the National Academy of Sciences 80, 1387–1391 (1983).

7 Jackson, J. B. C., Buss, L. W. & Cook, R. E. Population biology and evolution of clonal organisms. (Yale University Press, 1985).

8 Harper, J. L. Population Biology of Plants. (Academic Press, 1977).

9 Gaul, H. Die verschiedenen Bezugssysteme der Mutationshäufigkeit bei Pflanzen, angewendet auf Dosis-Effektkurven. Zeitschrift für Pflanzenzüchtung 38, 63–76 (1957).

10 Larkin, P. J. & Scowcroft, W. R. Somaclonal variation — a novel source of variability from cell cultures for plant improvement. Theoretical and Applied Genetics 60, 197–214 (1981).

11 Klekowski, E. J. & Kazarinovafukshansky, N. Shoot Apical Meristems and Mutation - Selective Loss of Disadvantageous Cell Genotypes. American Journal of Botany 71, 28–34 (1984).

12 Sutherland, W. J. & Watkinson, A. R. Somatic mutation: Do plants evolve differently? Nature 320, 305–305 (1986).

13 Fagerström, T., Briscoe, D. A. & Sunnucks, P. Evolution of mitotic cell-lineages in multicellular organisms. Trends in Ecology and Evolution 13, 117–120 (1998).

14 Lynch, M. Evolution of the mutation rate. Trends in Genetics 26, 345–352 (2010).

15 Gill, D. E., Chao, L., Perkins, S. L. & Wolf, J. B. Genetic mosaicism in plants and clonal animals. Annual Review in Ecology and Systematics 26, 423–444 (1995).

16 Antolin, M. F. & Strobeck, C. The population genetics of somatic mutations. The American Naturalist 126, 52–62 (1985).

17 Breese, E. L., Hayward, M. D. & Thomas, A. C. Somatic selection in perennial ryegrass. Heredity 20, 367 (1965).

18 Santelices, B., Gallegos Sánchez, C. & González, A. V. Intraorganismal genetic heterogeneity as a source of genetic variation in modular macroalgae. Journal of Phycology 54, 767–771 (2018).

19 Schoen, D. J. & Schultz, S. T. Somatic Mutation and Evolution in Plants. Annual Review of Ecology, Evolution, and Systematics in press, (2019).

20 Simberloff, D. & Leppanen, C. Plant somatic mutations in nature conferring insect and herbicide resistance. Pest Management Science 75, 14–17 (2019).

21 Reusch, T. B. H. & Boström, C. Widespread genetic mosaicism in the marine angiosperm Zostera marina is correlated with clonal reproduction. Evol Ecol 25, 899–913 (2011).

22 Arnaud-Haond, S. et al. Implications of Extreme Life Span in Clonal Organisms: Millenary Clones in Meadows of the Threatened Seagrass Posidonia oceanica. PLOS One 7, e30454 (2012).

23 Bricker, E., Calladine, A., Virnstein, R. & Waycott, M. Mega Clonality in an Aquatic Plant—A Potential Survival Strategy in a Changing Environment. Frontiers in Plant Science 9(2018).

24 Olsen, J. L. et al. The genome of the seagrass Zostera marina reveals angiosperm adaptation to the sea. Nature 530, 331–335 (2016).

25 Sung, W. et al. Evolution of the Insertion-Deletion Mutation Rate Across the Tree of Life. G3: Genes|Genomes|Genetics 6, 2583 (2016).

26 Poethig, S. Genetic mosaics and cell lineage analysis in plants. Trends in Genetics 5, 273–277 (1989).

27 Pineda-Krch, M. & Lehtilä, K. Cell Lineage Dynamics in Stratified Shoot Apical Meristems. Journal of Theoretical Biology 219, 495–505 (2002).

28 Klekowski, E. J. Plant clonality, mutation, diplontic selection and mutational meltdown. Biological Journal of the Linnean Society 79, 61–67 (2003).

29 Burian, A., Barbier de Reuille, P. & Kuhlemeier, C. Patterns of Stem Cell Divisions Contribute to Plant Longevity. Current Biology 26, 1385–1394 (2016).

30 Lang, G. I. et al. Pervasive genetic hitchhiking and clonal interference in forty evolving yeast populations. Nature 500, 571–574 (2013).

31 Williams, M. J., Werner, B., Barnes, C. P., Graham, T. A. & Sottoriva, A. Identification of neutral tumor evolution across cancer types. Nature Genetics 48, 238 (2016).

32 Williams, M. J. et al. Quantification of subclonal selection in cancer from bulk sequencing data. Nature Genetics 50, 895–903 (2018).

33 Schultz, Stewart T. & Scofield, Douglas G. Mutation Accumulation in Real Branches: Fitness Assays for Genomic Deleterious Mutation Rate and Effect in Large-Statured Plants. The American Naturalist 174, 163–175 (2009).

34 Willis, J. H. Inbreeding Load, Average Dominance and the Mutation Rate for Mildly Deleterious Alleles in Mimulus guttatus. Genetics 153, 1885–1898 (1999).

35 Otto, S. P. & Orive, M. E. Evolutionary consequences of mutation and selection within an individual. Genetics 141, 1173 (1995).

36 Orive, M. E. Somatic Mutations in Organisms with Complex Life Histories. Theoretical Population Biology 59, 235–249 (2001).

37 Otto, S. P. & Hastings, I. M. Mutation and selection within the individual. Genetica 102, 507 (1998).

38 Tarabichi, M. et al. Neutral tumor evolution? Nature Genetics 50, 1630–1633 (2018).

39 Frank, M. H. & Chitwood, D. H. Plant chimeras: The good, the bad, and the ‘Bizzaria’. Developmental Biology 419, 41–53 (2016).

40 Smith, M. L., Bruhn, J. N. & Anderson, J. B. The fungus Armillaria bulbosa is among the largest and oldest living organisms. Nature 356, 428–431 (1992).

41 Schmid-Siegert, E. et al. Low number of fixed somatic mutations in a long-lived oak tree. Nature Plants 3, 926–929 (2017).

42 Plomion, C. et al. Oak genome reveals facets of long lifespan. Nature Plants 4, 440–452 (2018).

43 de Witte, L. C. & Stöcklin, J. Longevity of clonal plants: why it matters and how to measure it. Annals of Botany 106, 859–870 (2010).

44 Ally, D., Ritland, K. & Otto, S. P. Aging in a Long-Lived Clonal Tree. PLOS Biology 8, e1000454 (2010).

45 Buss, L. W. The Evolution of Individuality. (Princeton University Press, 1987).

46 Santelices, B. How many kinds of individuals are there? Trends in Ecology and Evolution 14, 152–155 (1999).

47 Van Oppen, M. J. H., Souter, P., Howells, E. J., Heyward, A. & Berkelmans, R. Novel Genetic Diversity Through Somatic Mutations: Fuel for Adaptation of Reef Corals? Diversity 3, 405–423 (2011).

48 Gustafsson, C. & Boström, C. Algal mats reduce eelgrass (Zostera marina L.) growth in mixed and monospecific meadows. J Exp Mar Biol Ecol 461, 85–92 (2014).

49 Reusch, T. B. H., Chapman, A. R. O. & Gröger, J. P. Blue mussels (Mytilus edulis) do not interfere with eelgrass (Zostera marina) but fertilize shoot growth through biodeposition. Mar. Ecol. Prog. Ser. 108, 265–282 (1994).

50 Gustafsson, B. G. & Westman, P. On the causes for salinity variations in the Baltic Sea during the last 8500 years. Paleoceanography 17, 12-11-12-14 (2002).

51 Andrews, S. FastQC: a quality control tool for high throughput sequence data. (<https://www.bioinformatics.babraham.ac.uk/projects/fastqc/> 2010).

52 Bolger, A. M., Lohse, M. & Usadel, B. Trimmomatic: a flexible trimmer for Illumina sequence data. Bioinformatics 30, 2114–2120 (2014).

53 Li, H. & Durbin, R. Fast and accurate short read alignment with Burrows–Wheeler transform. Bioinformatics 25, 1754–1760 (2009).

54 Li, H. et al. The Sequence Alignment/Map format and SAMtools. Bioinformatics 25, 2078–2079 (2009).

55 Van der Auwera, G. A. et al. From FastQ Data to High-Confidence Variant Calls: The Genome Analysis Toolkit Best Practices Pipeline. Current Protocols in Bioinformatics 43, 11.10.11-11.10.33 (2013).

56 Ginestet, C. ggplot2: Elegant Graphics for Data Analysis. Journal of the Royal Statistical Society: Series A (Statistics in Society) 174, 245–246 (2011).

57 Danecek, P. et al. The variant call format and VCFtools. Bioinformatics 27, 2156–2158 (2011).

58 Noorbakhsh, J. & Chuang, J. H. Uncertainties in tumor allele frequencies limit power to infer evolutionary pressures. Nature Genetics 49, 1288 (2017).

59 Cingolani, P. et al. A program for annotating and predicting the effects of single nucleotide polymorphisms, SnpEff. Fly 6, 80–92 (2012).

60 Paradis, E. & Schliep, K. ape 5.0: an environment for modern phylogenetics and evolutionary analyses in R. Bioinformatics 35, 526–528 (2018).

61 Core Team, R. R: A language and environment for statistical computing. (R Foundation for Statistical Computing, Vienna, Austria, URL https://www.r-project.org/. 2019).

62 Kamvar, Z. N., Brooks, J. C. & Grünwald, N. J. Novel R tools for analysis of genome-wide population genetic data with emphasis on clonality. Frontiers in Genetics 6, 208 (2015).

63 Abyzov, A., Urban, A. E., Snyder, M. & Gerstein, M. CNVnator: An approach to discover, genotype, and characterize typical and atypical CNVs from family and population genome sequencing. Genome Research 21, 974–984 (2011).

64 Robinson, J. T., Thorvaldsdóttir, H., Wenger, A. M., Zehir, A. & Mesirov, J. P. Variant Review with the Integrative Genomics Viewer. Cancer Research 77, e31 (2017).

65 Quinlan, A. R. & Hall, I. M. BEDTools: a flexible suite of utilities for comparing genomic features. Bioinformatics 26, 841–842 (2010).

66 Cibulskis, K. et al. Sensitive detection of somatic point mutations in impure and heterogeneous cancer samples. Nature Biotechnology 31, 213 (2013).

67 Bataillon, T. et al. Inference of Purifying and Positive Selection in Three Subspecies of Chimpanzees (Pan troglodytes) from Exome Sequencing. Genome Biology and Evolution 7, 1122–1132 (2015).

68 Alexa, A. & Rahnenfuhrer, J. topGO: Enrichment Analysis for Gene Ontology. (R package version 2.36.0.a, 2019).

