## Supplementary_Figs_Tables_all for "Somatic genetic drift and multi-level selection in modular species"

#### Supplementary information

Supplementary Dataset 1: Detailed information for 432 fixed non-synonymous SNP

**Gelöscht:** s

#### Supplementary Figures

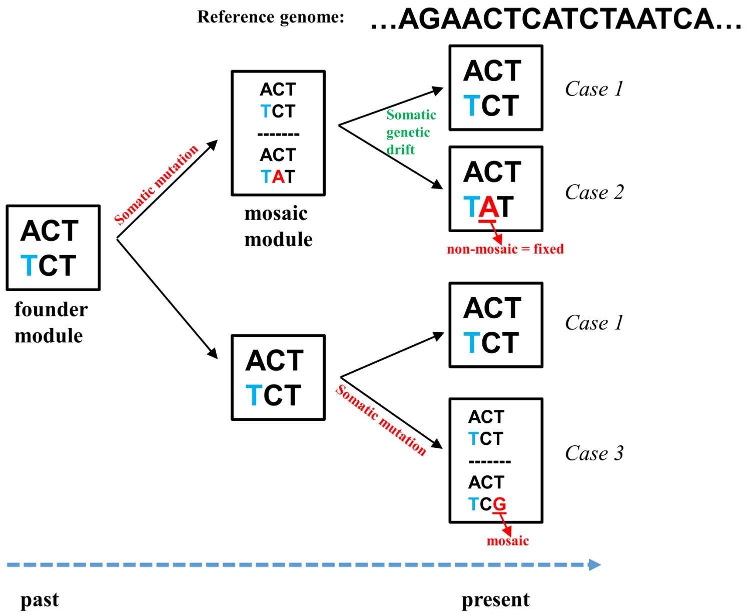

**Supplementary Figure 1: Somatic genetic drift in modular organisms and possible origins and categories of single nucleotide polymorphisms (SNPs).** Multicellular organisms originate from a single zygote and achieve growth and development via mitosis. The diagram indicates possible genotypes at a locus within the module (=box). Heterozygous polymorphisms present in the zygote (blue) are passed on to all the offspring cells, and result in consistent heterozygous calls across all modules. During growth, cells acquire somatic mutations (red) owing to mitotic errors that initially emerge as genetic mosaics, i.e. the somatic polymorphisms are present in only a subset of cells. When modular organisms form another iterative unit (=module), a few cells in the parent module are progenitors for the new module. The bottleneck effects accompanying this random sampling process determine the somatic polymorphisms in the new modules through somatic genetic drift. Somatic genetic drift thus converts mosaic somatic polymorphisms into different fixed module genotypes. In case 1, somatic genetic drift restores the wild-type genotype present in the zygote. In cases 2, somatic genetic drift segregates the mosaic genetic diversity at that locus into a fixed state. In case 3, a novel mutation emerges in a mosaic state, which would require additional rounds of somatic genetic drift to either being fixed or lost.

### Inter- and intra-module SNP calling workflow

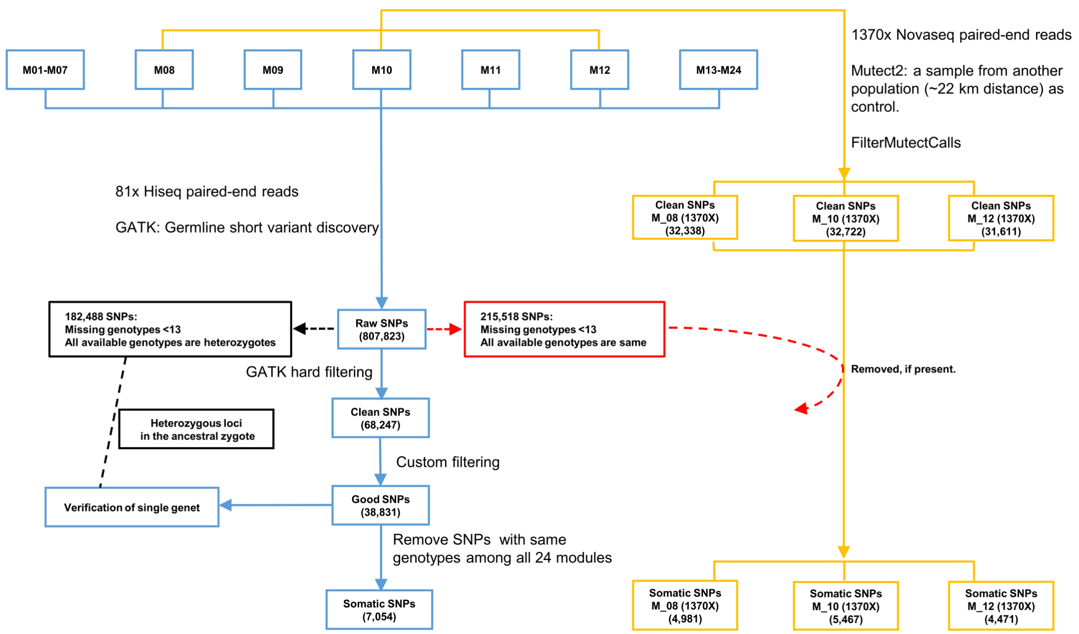

**Supplementary Figure 2: Workflow for inter- and intra-module SNP calling.** All 24 modules were sequenced at an average coverage of 81x on Hiseq 4000 platform, and a subset of three modules (M08, M10, and M12) were further sequenced at an average coverage of 1370x on Novaseq 6000 platform. The 81x dataset was used to call fixed (=fully heterozygous) SNPs among modules, estimate the number of heterozygous loci in the initial zygote, and locate SNPs that were genetically different between the reference genome and the initial zygote. The 1370x dataset was used to call intra-module somatic polymorphisms, and to verify the proportion of mosaic vs. fully heterozygous SNPs.

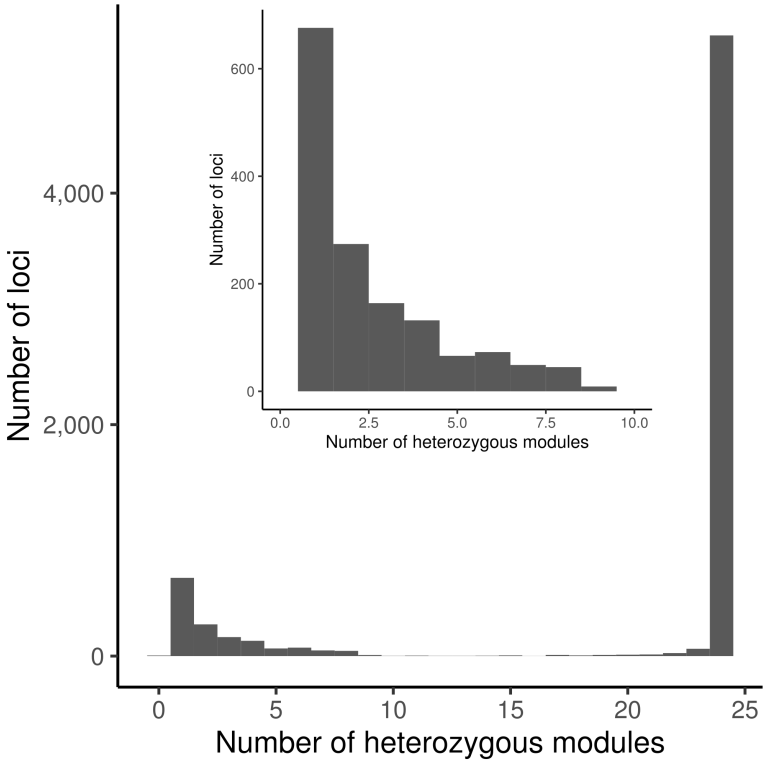

**Supplementary Figure 3: Verification of modules belonging to a single genet of *Z. marina* based on indels.** The fully heterozygous indel polymorphisms among the 24 modules were also used to verify if they belong to the same genet. The result is very similar to that based on fixed SNPs (Fig. 2b). The x-axis indicates the number of modules that are heterozygous at a given locus, and the y axis indicates the number of loci. Under exclusively asexual reproduction, the heterozygous loci in the initial zygote will be passed on to all subsequent cells and modules, and result in a single dominant peak indicating the loci where all modules are identically heterozygous.

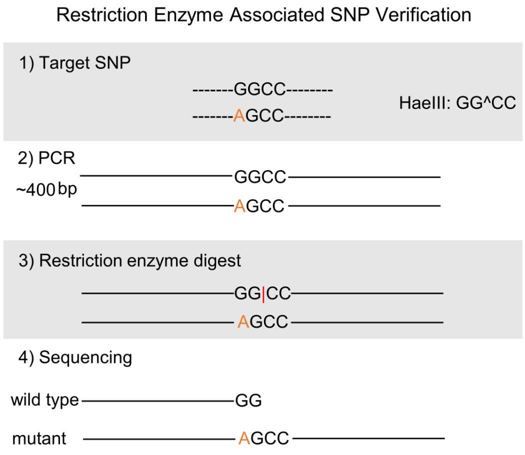

**Supplementary Figure 4: Restriction-enzyme based SNP verification method.** In order to verify SNP loci independent of any sequencing technology, we first searched the reference genome for restriction enzyme motifs covering the target SNPs. Along with the reference genome, our target genotype possesses only intact motif when homozygous that will be recognized by a selected restriction enzyme, while the variant allele changes the recognition sequence. Fluorescence-labelled primer pairs were designed to amplify the ~400 bp amplicon encompassing the target SNP, which was subsequently digested with an appropriate restriction enzyme. Heterozygous samples display fluorescent peaks at two different positions, while the homozygous genotype has only one signal (i.e. both homologous sites are cut at the same position).

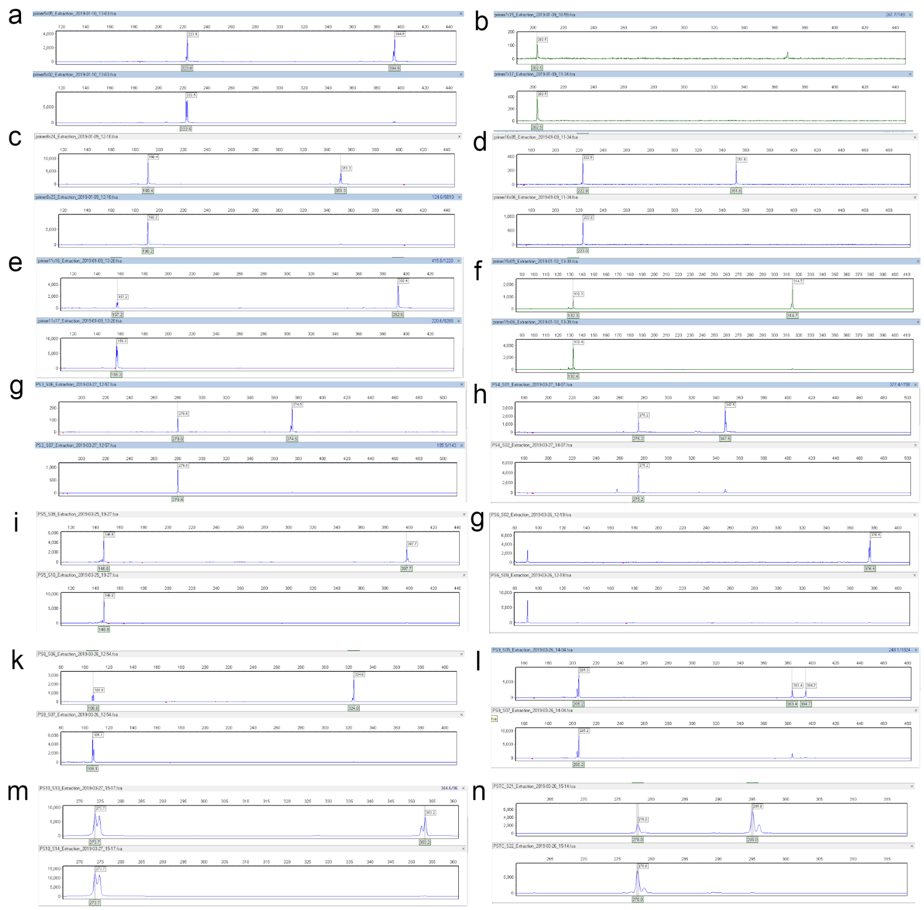

**Supplementary Figure 5: Experimental verification of fixed SNPs among *Z. marina* modules.** In all panels (a-n), the upper graph with fluorescent peaks at two different positions represents the heterozygous genotype. The panels (a-n) indicate the following loci: 1st_05, 1st_07, 1st_08, 1st_10, 1st_11, 1st_15, 2nd_03, 2nd_04, 2nd_05, 2nd_06, 2nd_08, 2nd_09, 2nd_10, and 2nd_TC. The detailed information for each locus can be found in Supplementary Table 4. In some cases, such as panel (m), the fluorescent signal strength was too high (>10,000) which led to a twin peak, which however does not interfere with a proper interpretation. All 14 loci in combination with 24 modules were confirmed.

**Gelöscht:** 8

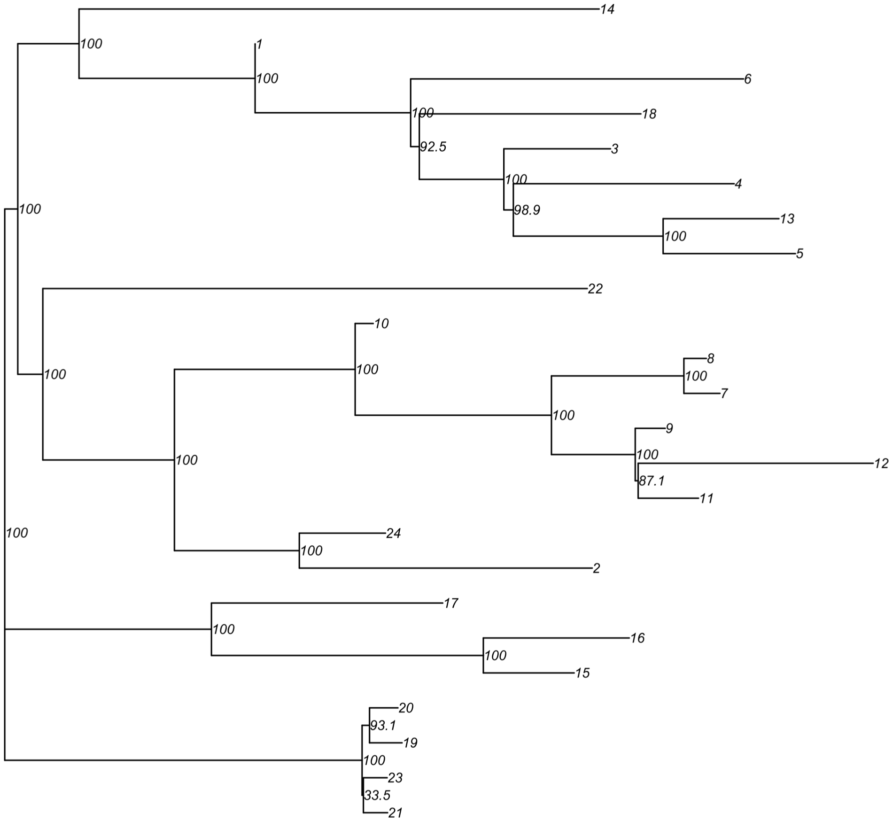

**Supplementary Figure 6: Bootstrap values accompanying the NJ tree depicted in Figure 2d.**

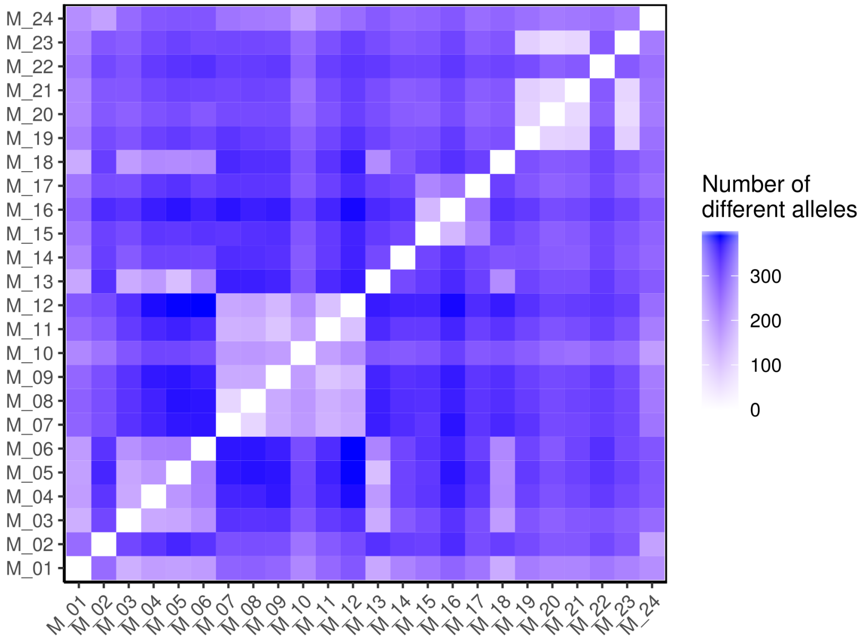

**Supplementary Figure 7: Pairwise genetic differences among 24 seagrass *Z. marina* modules based on small insertion-deletion polymorphisms (indels).** 1,654 fixed indels among the 24 modules were used to calculate pairwise genetic differences based on the number of different alleles. The result is consistent to the heat map based on fully heterozygous (=fixed) SNPs (Fig. 2c).

Gelöscht: s

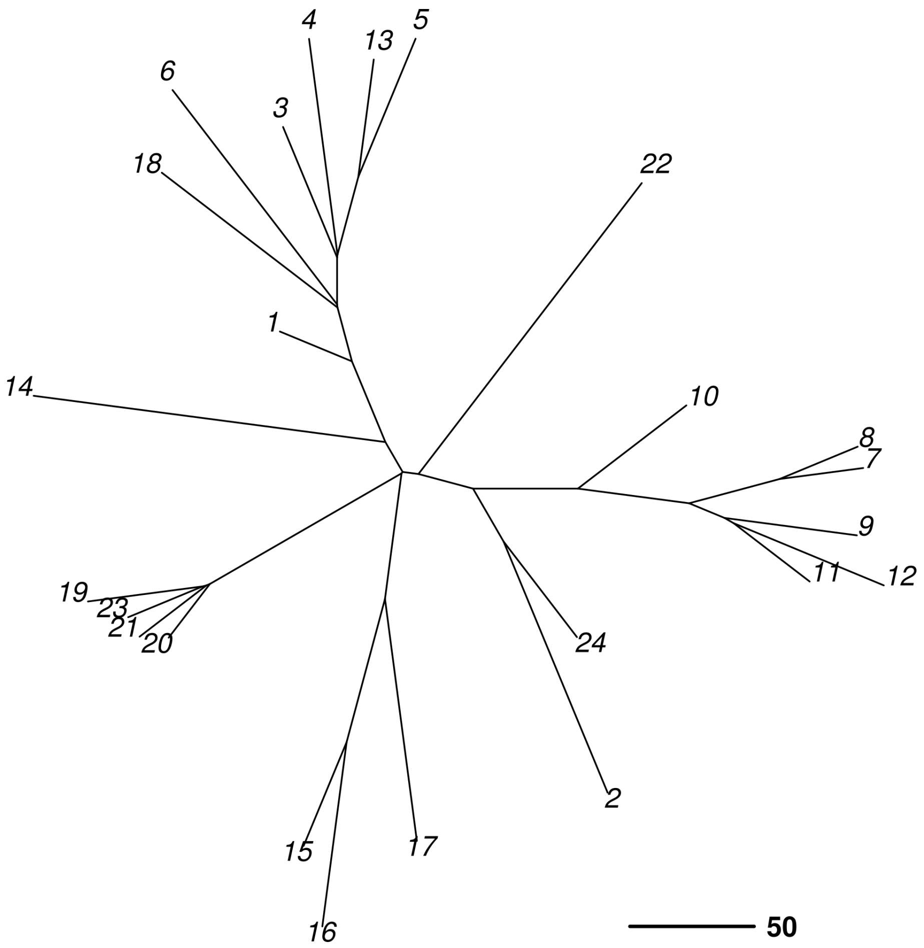

**Supplementary Figure 8: Neighbor-joining tree for 24 seagrass *Z. marina* modules based on indels.** The 1,654 fixed indel polymorphisms among the 24 modules were used to quantify the pairwise genetic differences, based on the calculated number of different alleles. The genetic distance matrix was used to construct a neighbor-joining tree, which is similar to that revealed by fixed SNPs (Fig. 2d).

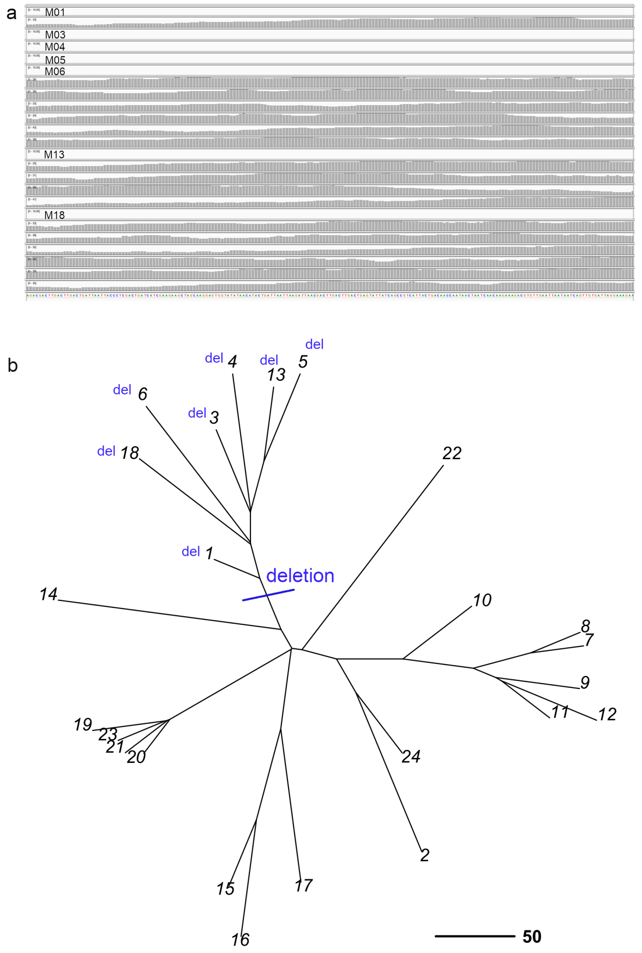

**Supplementary Figure 9: A large structural polymorphism among *Z. marina* modules.** We found a >200 bp structural polymorphism among the 24 modules. Its presence/absence among the 24 modules is consistent with the topology depicted by the NJ tree.

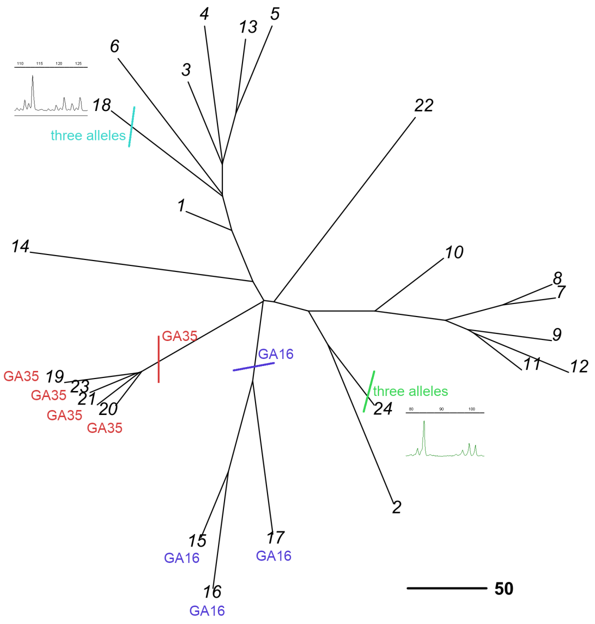

**Supplementary Figure 10: Microsatellite polymorphisms among 24 seagrass *Z. marina* modules.** We genotyped all modules for 8 microsatellite loci (Genbank accession numbers: AJ009904.1, AJ009901.1, AJ249304.1, AJ249306.1, AJ009898.1, AJ009900.1, AJ249307.1, and AJ249305.1) and detected 7 somatic polymorphisms. Their presence/absence among the 24 modules is consistent with the NJ tree. Except for the clear mosaic ones, these have been interpreted as segregating genetic variation without the SNP calling based on full genomes (Fig. 2b). Five microsatellite alleles were only present in one module, two of which appeared as a third, mosaic allele. Two polymorphisms were present in more than one modules. Small insets show the three-allelic, mosaic genotypes.

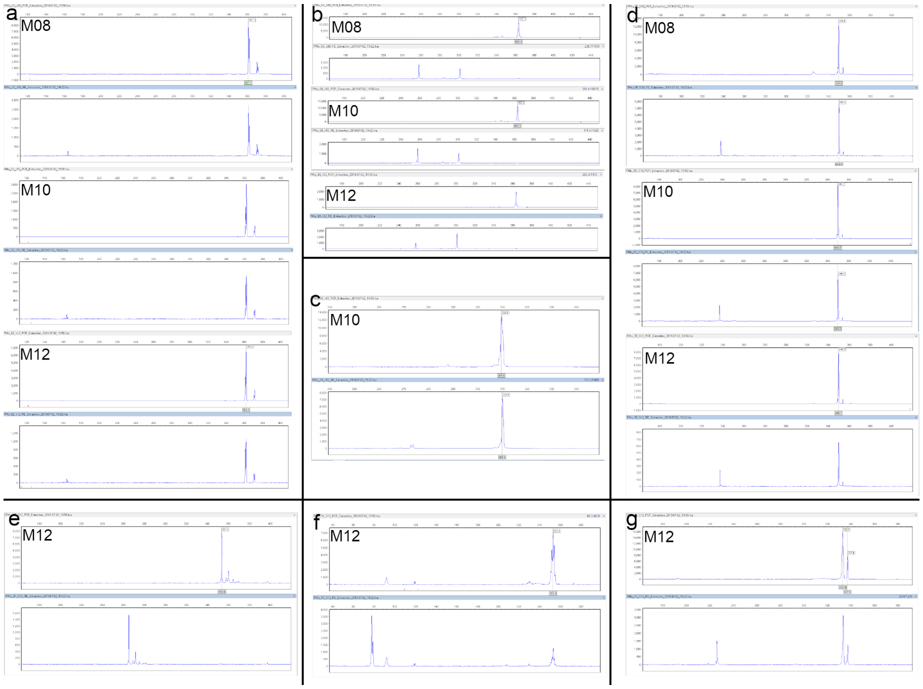

**Supplementary Figure 11: Verification of mosaic, intra-module somatic polymorphisms in *Z. marina*.** We verified mosaic SNPs in that the variant allele converted a non-restriction site to a restriction site. Thus, after restriction enzyme digestion shorter PCR fragments will appear only if the variant allele is present, an observation that cannot be biased by undigested wild-type alleles. For each combination of locus and sample (e.g. a + M08), the upper graph and the lower graph represent the PCR products before and after restriction enzyme digest, respectively. The variant allele is represented as a shorter fragment in the lower graph. Panels (a-g) represent the following loci: Mu_02, Mu_03, Mu_04, Mu_05, Mu_07, Mu_11, and Mu_12. The detailed information for each locus can be found in Supplementary Table 8. In panel (b), the amplicon contains another restriction site ("GGCC") at the downstream of the target SNP, so all the amplicons were universally cleaved into a shorter one, based on which the restriction enzyme cut another time on the target site.

**Gelöscht:** 9

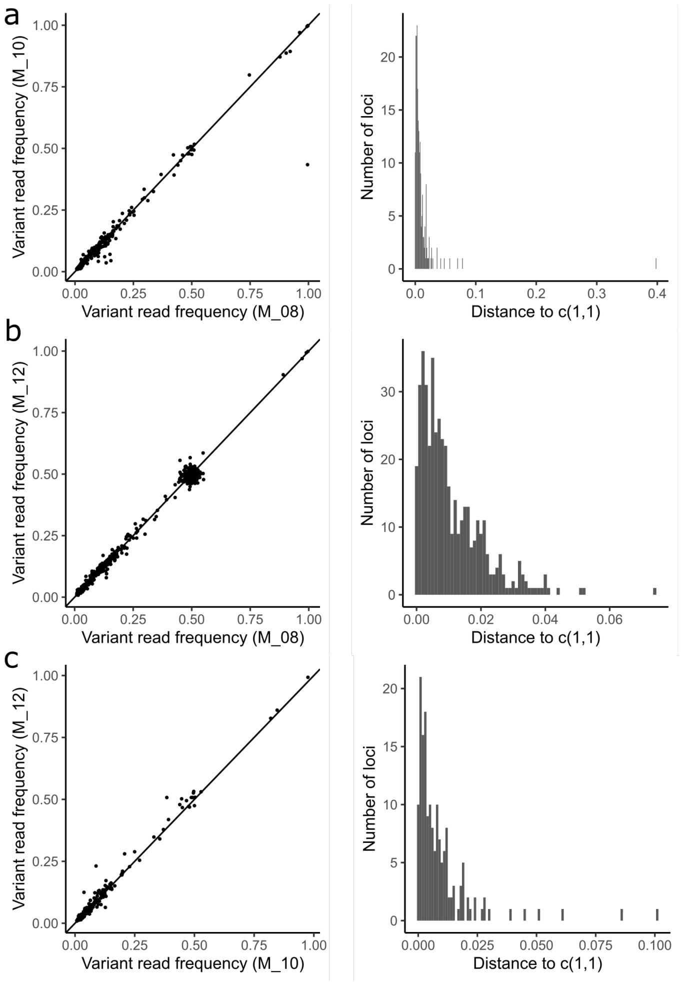

**Supplementary Figure 12: The intra-module somatic polymorphisms shared by two modules show stable allele frequencies.** At the intra-module section (1370x dataset, modules M08, M10, and M12), the somatic polymorphisms were categorized into 1) being present in only one module, 2) being shared by two modules, and 3) being shared by all three modules (cf. Venn diagram Fic 3a). Plots are based on SNPs shared by two modules, in panel **a** among M08 and M10, in **b** among M08 and M12, in **c** among M10 and M12. The frequencies of a somatic polymorphism in the two modules determine its position in the two-dimension coordinate system. Most somatic polymorphisms are all close to the line y=x, showing stable frequencies. The distance of a position to the line y=x indicates the difference of allele frequency in the two modules shown in a histogram on the right of each panel.

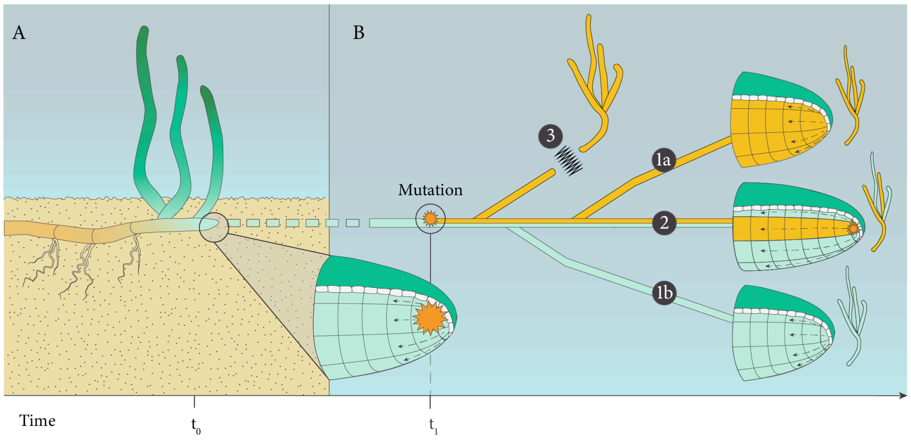

**Supplementary Figure 13: *Z. marina* modules exist as physiologically independent and genetically unique "individuals".** Somatic mutations emerge in a mosaic state. Along with the growth of the plant, somatic genetic drift segregates the mosaic genetic diversity into different fixed genotypes among modules (case 1a & 1b). In other modules, they may remain a mosaic state as well (case 2). Note that the modules are physiologically independent, as the intermediate rhizome connecting the modules will rot away within one or two years (case 3).

#### Supplementary Tables

**Supplementary Table 1: Sampling design.** Twenty-four modules of the seagrass *Z. marina* were collected along a predetermined inshore-offshore transect consisting of two sections, one at ~2 m depth and the other one at ~4 m depth. Twelve modules were collected in each section, respectively.

| SHALLOW | | | DEEP | | |
|---|---|---|---|---|---|
| **Sample** | **Depth (m)** | **Distance from starting point (m)** | **Sample** | **Depth (m)** | **Distance from starting point (m)** |
| M_01 | 2.1 | 0 | M_13 | 4.1 | 0 |
| M_02 | 2.4 | 10 | M_14 | 3.9 | 10 |
| M_03 | 2.4 | 20 | M_15 | 3.8 | 20 |
| M_04 | 2.0 | 30 | M_16 | 4.1 | 30 |
| M_05 | 2.0 | 44 | M_17 | 4.2 | 40 |
| M_06 | 2.3 | 54 | M_18 | 4.4 | 55 |
| M_07 | 2.4 | 65 | M_19 | 4.4 | 68 |
| M_08 | 2.4 | 75 | M_20 | 4.5 | 78 |
| M_09 | 1.9 | 88 | M_21 | 4.5 | 89 |
| M_10 | 2.3 | 100 | M_22 | 4.6 | 99 |
| M_11 | 2.5 | 111 | M_23 | 4.8 | 109 |
| M_12 | 2.6 | 120 | M_24 | 5.1 | 119 |

**Supplementary Table 2: Basic statistics on whole-genome resequencing of 24 *Z. marina* modules on an Illumina Hiseq 4000 platform.** All 24 modules were sequenced in a paired-end (2 × 151bp) mode to 81× coverage on average.

| Sample | Read | Raw reads (bases) | Trimmed reads (bases) | Trimmed reads (coverage) | Mapping rate (%) |
|---|---|---|---|---|---|
| M_01 | R1 | 6,514,773,143 | 4,784,239,699 | 48.05 | 93.45 |
| | R2 | 6,514,773,143 | 5,012,644,377 | | |
| M_02 | R1 | 9,126,143,285 | 8,239,834,760 | 76.87 | 94.90 |
| | R2 | 9,126,143,285 | 7,435,005,872 | | |
| M_03 | R1 | 10,248,138,970 | 9,146,298,916 | 85.69 | 95.01 |
| | R2 | 10,248,138,970 | 8,327,087,256 | | |
| M_04 | R1 | 8,712,513,364 | 7,797,060,447 | 72.56 | 92.80 |
| | R2 | 8,712,513,364 | 6,999,344,576 | | |
| M_05 | R1 | 12,247,961,075 | 9,794,025,318 | 94.53 | 94.37 |
| | R2 | 12,247,961,075 | 9,480,772,227 | | |
| M_06 | R1 | 10,066,487,933 | 9,104,901,065 | 84.89 | 95.48 |
| | R2 | 10,066,487,933 | 8,205,080,108 | | |
| M_07 | R1 | 10,018,128,975 | 9,114,359,843 | 85.37 | 94.30 |
| | R2 | 10,018,128,975 | 8,293,245,178 | | |
| M_08 | R1 | 10,666,580,808 | 9,713,578,994 | 90.71 | 93.65 |
| | R2 | 10,666,580,808 | 8,782,113,457 | | |
| M_09 | R1 | 9,414,017,839 | 8,582,514,949 | 80.60 | 92.59 |
| | R2 | 9,414,017,839 | 7,851,937,650 | | |
| M_10 | R1 | 6,528,798,325 | 5,940,882,792 | 55.49 | 94.98 |
| | R2 | 6,528,798,325 | 5,373,273,275 | | |
| M_11 | R1 | 9,783,097,626 | 8,990,162,245 | 84.53 | 94.84 |
| | R2 | 9,783,097,626 | 8,246,472,161 | | |
| M_12 | R1 | 11,572,405,799 | 10,554,499,766 | 98.95 | 95.24 |
| | R2 | 11,572,405,799 | 9,622,152,935 | | |

| | | | | | |
|---|---|---|---|---|---|
| M_13 | R1 | 9,083,441,089 | 8,283,428,213 | 77.78 | 93.34 |
| | R2 | 9,083,441,089 | 7,576,753,589 | | |
| M_14 | R1 | 10,169,361,968 | 9,277,920,776 | 87.14 | 94.09 |
| | R2 | 10,169,361,968 | 8,489,958,466 | | |
| M_15 | R1 | 9,313,684,228 | 8,483,003,078 | 79.84 | 94.99 |
| | R2 | 9,313,684,228 | 7,798,061,512 | | |
| M_16 | R1 | 10,977,353,002 | 10,050,369,523 | 94.66 | 92.11 |
| | R2 | 10,977,353,002 | 9,252,291,537 | | |
| M_17 | R1 | 11,060,730,068 | 10,028,027,166 | 93.81 | 94.17 |
| | R2 | 11,060,730,068 | 9,101,583,955 | | |
| M_18 | R1 | 7,425,409,296 | 6,727,127,750 | 63.06 | 93.98 |
| | R2 | 7,425,409,296 | 6,132,131,423 | | |
| M_19 | R1 | 9,126,426,108 | 8,363,753,419 | 78.78 | 92.75 |
| | R2 | 9,126,426,108 | 7,699,515,554 | | |
| M_20 | R1 | 10,223,403,509 | 8,959,008,457 | 85.95 | 94.09 |
| | R2 | 10,223,403,509 | 8,566,982,853 | | |
| M_21 | R1 | 9,069,893,671 | 8,321,208,742 | 78.74 | 93.77 |
| | R2 | 9,069,893,671 | 7,733,756,450 | | |
| M_22 | R1 | 9,349,276,438 | 8,484,251,652 | 79.70 | 93.44 |
| | R2 | 9,349,276,438 | 7,767,904,874 | | |
| M_23 | R1 | 9,959,155,472 | 9,137,826,475 | 86.27 | 93.31 |
| | R2 | 9,959,155,472 | 8,453,672,321 | | |
| M_24 | R1 | 10,101,766,969 | 9,250,283,727 | 87.08 | 94.74 |
| | R2 | 10,101,766,969 | 8,506,391,862 | | |

**Supplementary Table 3: NCBI accession numbers.** The raw reads were submitted to SRA database.

| Sample | BioSample accession number | BioProject accession number |
|---|---|---|
| M_01 | SAMN12389300 | PRJNA557092 |
| M_02 | SAMN12389301 | PRJNA557092 |
| M_03 | SAMN12389302 | PRJNA557092 |
| M_04 | SAMN12389303 | PRJNA557092 |
| M_05 | SAMN12389304 | PRJNA557092 |
| M_06 | SAMN12389305 | PRJNA557092 |
| M_07 | SAMN12389306 | PRJNA557092 |
| M_08 | SAMN12389307 | PRJNA557092 |
| M_09 | SAMN12389308 | PRJNA557092 |
| M_10 | SAMN12389309 | PRJNA557092 |
| M_11 | SAMN12389310 | PRJNA557092 |
| M_12 | SAMN12389311 | PRJNA557092 |
| M_13 | SAMN12389312 | PRJNA557092 |
| M_14 | SAMN12389313 | PRJNA557092 |
| M_15 | SAMN12389314 | PRJNA557092 |
| M_16 | SAMN12389315 | PRJNA557092 |
| M_17 | SAMN12389316 | PRJNA557092 |
| M_18 | SAMN12389317 | PRJNA557092 |
| M_19 | SAMN12389318 | PRJNA557092 |
| M_20 | SAMN12389319 | PRJNA557092 |
| M_21 | SAMN12389320 | PRJNA557092 |
| M_22 | SAMN12389321 | PRJNA557092 |
| M_23 | SAMN12389322 | PRJNA557092 |
| M_24 | SAMN12389323 | PRJNA557092 |

**Supplementary Table 4: Detailed information for 14 verified fixed SNPs.** Fully heterozygous SNPs of 14 loci were verified across all 24 modules. Example electropherograms of verification results can be found in Supplementary Figure 5.

| Locus | Contig No. | Position | Enzyme | Restriction site | 5' Fluorescent dye | Forward primer | Reverse primer |
|---|---|---|---|---|---|---|---|
| 1st_05 | LFYR01001079.1 | 70,478 | BamHI | G^GATC_C | FAM | AGGTGGTTTGACTTCCGTCC | ACCATGCAAGAGCCCCTAAC |
| 1st_07 | LFYR01000619.1 | 331,575 | HaeIII | GG^_CC | HEX | AGGGGTTTTTGTTCGGTCGA | TTTCAGTCGAGACAGGGTGC |
| 1st_08 | LFYR01000671.1 | 881,966 | HaeIII | GG^_CC | FAM | TCCAGAACCAGCATTGACGA | AAAGATGCAGCCAGGGGAAG |
| 1st_10 | LFYR01001213.1 | 1,012,778 | HaeIII | GG^_CC | FAM | TGATCTTTGGTCGTGGAGCA | TCGGGAGGTTGGTCTCTCAT |
| 1st_11 | LFYR01001351.1 | 610,127 | HaeIII | GG^_CC | FAM | AAGTCCATCAAGAGCACGCA | CGTCGCCGGATCCTATGAAA |
| 1st_15 | LFYR01001054.1 | 847,248 | NcoI | C^CATG_G | HEX | GTGTGGGTTCGCCAGAGTTA | GCGTTTGTGAAATGGCGACT |
| 2nd_03 | LFYR01000655.1 | 208,328 | HaeIII | GG^_CC | FAM | TGGTTCGCAGCATGAATCCT | GCAGTCAAGAGGTGTGGTGT |
| 2nd_04 | LFYR01000709.1 | 341,678 | HaeIII | GG^_CC | FAM | TTGGATTGAGCCCGATGTCC | CGCTTTATGCTGGACCCTGA |
| 2nd_05 | LFYR01000781.1 | 174,973 | HaeIII | GG^_CC | FAM | CCCCTTTGTTGGATCTTCTTCC | TCGGCTCGGCTTGATAAACT |
| 2nd_06 | LFYR01000915.1 | 494,331 | HaeIII | GG^_CC | FAM | TGACTTGGAACCTCCCCAGA | ACGGCGCATTTCTGTCAAAA |
| 2nd_08 | LFYR01001661.1 | 436,971 | HaeIII | GG^_CC | FAM | ACACCAAGAGCAATCAACCT | CACATTCCAGGCGCAAAACA |
| 2nd_09 | LFYR01001977.1 | 179,811 | HaeIII | GG^_CC | FAM | TTTCAGCAAAGGCAGCCAAG | GCCAGCCCTTCCTCTTTGAT |
| 2nd_10 | LFYR01002125.1 | 147,686 | HaeIII | GG^_CC | FAM | CGTAGGTTTGCCTCGTCACT | CGCTCCATAGTCTGCCGATT |
| 2nd_TC | LFYR01002184.1 | 54,937 | HaeIII | GG^_CC | FAM | ACAACGCTAGGAGACATGTTCT | CGATTCTACATTACCGGCCCA |

**Gelöscht:** ea
**Gelöscht:** ea
**Gelöscht:** ea
**Gelöscht:** e
**Gelöscht:** a
**Gelöscht:** ea
**Gelöscht:** ea
**Gelöscht:** ea
**Gelöscht:** ea

**Supplementary Table 5: Pairwise genetic distance matrix.** The genetic distance was quantified by number of different alleles. The upper half was calculated based on 7,054 fixed (=fully heterozygous) SNPs, and the lower half was calculated based on 432 fixed nonsynonymous SNPs.

| | **M_01** | **M_02** | **M_03** | **M_04** | **M_05** | **M_06** | **M_07** | **M_08** | **M_09** | **M_10** | **M_11** | **M_12** | **M_13** | **M_14** | **M_15** | **M_16** | **M_17** | **M_18** | **M_19** | **M_20** | **M_21** | **M_22** | **M_23** | **M_24** |
|---|---|---|---|---|---|---|---|---|---|---|---|---|---|---|---|---|---|---|---|---|---|---|---|---|
| **M_01** | - | 988 | 452 | 583 | 665 | 594 | 1,134 | 1,118 | 1,071 | 757 | 1,092 | 1,291 | 642 | 850 | 996 | 1,049 | 840 | 481 | 807 | 795 | 784 | 979 | 783 | 740 |
| **M_02** | 55 | - | 1,446 | 1,588 | 1,664 | 1,610 | 1,185 | 1,171 | 1,117 | 761 | 1,149 | 1,362 | 1,649 | 1,432 | 1,429 | 1,487 | 1,257 | 1,472 | 1,208 | 1,211 | 1,200 | 1,343 | 1,188 | 466 |
| **M_03** | 31 | 88 | - | 413 | 490 | 649 | 1,588 | 1,583 | 1,535 | 1,162 | 1,575 | 1,799 | 471 | 1,288 | 1,442 | 1,523 | 1,289 | 507 | 1,240 | 1,224 | 1,220 | 1,436 | 1,220 | 1,189 |
| **M_04** | 32 | 88 | 24 | - | 611 | 804 | 1,756 | 1,731 | 1,678 | 1,315 | 1,724 | 1,946 | 605 | 1,456 | 1,604 | 1,668 | 1,439 | 662 | 1,377 | 1,376 | 1,377 | 1,593 | 1,374 | 1,341 |
| **M_05** | 44 | 100 | 36 | 36 | - | 873 | 1,838 | 1,820 | 1,764 | 1,408 | 1,807 | 2,020 | 305 | 1,522 | 1,670 | 1,740 | 1,512 | 727 | 1,465 | 1,459 | 1,452 | 1,669 | 1,445 | 1,420 |
| **M_06** | 34 | 90 | 44 | 44 | 56 | - | 1,776 | 1,763 | 1,695 | 1,333 | 1,747 | 1,956 | 861 | 1,462 | 1,611 | 1,683 | 1,440 | 697 | 1,390 | 1,399 | 1,387 | 1,605 | 1,383 | 1,356 |
| **M_07** | 83 | 81 | 118 | 123 | 135 | 125 | - | 72 | 345 | 464 | 391 | 609 | 1,816 | 1,572 | 1,578 | 1,657 | 1,427 | 1,645 | 1,376 | 1,353 | 1,346 | 1,493 | 1,349 | 926 |
| **M_08** | 83 | 81 | 118 | 123 | 135 | 125 | 0 | - | 321 | 447 | 374 | 596 | 1,792 | 1,559 | 1,571 | 1,640 | 1,403 | 1,627 | 1,353 | 1,350 | 1,326 | 1,472 | 1,326 | 914 |
| **M_09** | 78 | 70 | 112 | 112 | 124 | 114 | 27 | 27 | - | 412 | 114 | 330 | 1,734 | 1,521 | 1,505 | 1,582 | 1,351 | 1,573 | 1,292 | 1,295 | 1,287 | 1,441 | 1,283 | 855 |
| **M_10** | 48 | 45 | 82 | 87 | 99 | 89 | 36 | 36 | 43 | - | 450 | 663 | 1,380 | 1,143 | 1,149 | 1,217 | 996 | 1,212 | 938 | 937 | 913 | 1,070 | 925 | 508 |
| **M_11** | 82 | 75 | 117 | 117 | 129 | 119 | 32 | 32 | 5 | 48 | - | 363 | 1,773 | 1,562 | 1,550 | 1,624 | 1,387 | 1,612 | 1,344 | 1,338 | 1,326 | 1,486 | 1,325 | 892 |
| **M_12** | 99 | 91 | 133 | 133 | 145 | 135 | 48 | 48 | 21 | 64 | 26 | - | 1,995 | 1,781 | 1,772 | 1,833 | 1,604 | 1,822 | 1,558 | 1,554 | 1,537 | 1,699 | 1,543 | 1,103 |
| **M_13** | 39 | 96 | 32 | 32 | 22 | 52 | 131 | 131 | 120 | 95 | 125 | 141 | - | 1,506 | 1,654 | 1,721 | 1,496 | 718 | 1,446 | 1,443 | 1,429 | 1,644 | 1,423 | 1,399 |
| **M_14** | 50 | 84 | 83 | 88 | 100 | 90 | 101 | 101 | 108 | 65 | 113 | 129 | 96 | - | 1,436 | 1,505 | 1,268 | 1,339 | 1,224 | 1,218 | 1,189 | 1,410 | 1,200 | 1,171 |
| **M_15** | 61 | 81 | 93 | 93 | 105 | 95 | 116 | 116 | 105 | 80 | 110 | 126 | 101 | 89 | - | 291 | 735 | 1,479 | 1,187 | 1,177 | 1,176 | 1,424 | 1,176 | 1,167 |
| **M_16** | 60 | 81 | 93 | 93 | 105 | 95 | 116 | 116 | 105 | 80 | 110 | 126 | 99 | 89 | 12 | - | 793 | 1,543 | 1,258 | 1,255 | 1,242 | 1,484 | 1,239 | 1,234 |
| **M_17** | 57 | 76 | 88 | 88 | 100 | 90 | 111 | 111 | 100 | 75 | 105 | 121 | 96 | 84 | 43 | 43 | - | 1,326 | 1,023 | 1,024 | 1,008 | 1,258 | 1,003 | 996 |
| **M_18** | 33 | 88 | 39 | 39 | 51 | 44 | 123 | 123 | 112 | 87 | 117 | 133 | 47 | 88 | 93 | 93 | 88 | - | 1,271 | 1,255 | 1,249 | 1,469 | 1,251 | 1,219 |
| **M_19** | 48 | 68 | 80 | 80 | 92 | 82 | 103 | 103 | 92 | 67 | 97 | 113 | 88 | 76 | 71 | 71 | 66 | 80 | - | 75 | 81 | 1,212 | 81 | 947 |
| **M_20** | 48 | 68 | 80 | 80 | 92 | 82 | 103 | 103 | 92 | 67 | 97 | 113 | 88 | 76 | 71 | 71 | 66 | 80 | 0 | - | 72 | 1,212 | 77 | 937 |
| **M_21** | 49 | 74 | 81 | 86 | 98 | 88 | 94 | 94 | 98 | 58 | 103 | 119 | 94 | 67 | 77 | 77 | 72 | 86 | 6 | 6 | - | 1,176 | 59 | 932 |
| **M_22** | 61 | 86 | 95 | 102 | 114 | 104 | 103 | 103 | 110 | 67 | 115 | 131 | 110 | 80 | 95 | 95 | 90 | 102 | 82 | 82 | 73 | - | 1,172 | 1,098 |
| **M_23** | 48 | 71 | 81 | 83 | 95 | 85 | 100 | 100 | 95 | 64 | 100 | 116 | 91 | 73 | 74 | 74 | 69 | 83 | 3 | 3 | 3 | 79 | - | 939 |
| **M_24** | 45 | 19 | 77 | 77 | 89 | 79 | 70 | 70 | 59 | 34 | 64 | 80 | 85 | 73 | 70 | 70 | 65 | 77 | 57 | 57 | 63 | 75 | 60 | - |

**Supplementary Table 6: Ultra-deep whole-genome resequencing of 3 seagrass *Z. marina* modules on an Illumina Novaseq 6000 platform.** Module_08, Module_10 and Module_12 were further sequenced on Novaseq 6000 platform at 1370× coverage on average (paired-end 2 × 151bp). Basic information and statistics are given.

| Sample | Read | Raw_reads (bases) | Trimmed_reads (bases) | Trimmed_reads (coverage) | Mapping rate (%) |
|---|---|---|---|---|---|
| M_08 | R1 | 133,813,282,831 | 126,375,440,334 | 1,238 | 93.63 |
| | R2 | 133,813,282,831 | 126,134,322,303 | | |
| M_10 | R1 | 139,403,148,056 | 131,822,147,904 | 1,295 | 95.01 |
| | R2 | 139,403,148,056 | 132,180,830,941 | | |
| M_12 | R1 | 170,779,135,150 | 160,947,684,601 | 1,578 | 95.19 |
| | R2 | 170,779,135,150 | 160,737,274,523 | | |

**Supplementary Table 7: Verification results for 9 low-frequency intra-module SNPs.** Mosaic SNPs were verified in one to three modules, depending on the SNP calling results. Note that both non-verified SNPs had very low variant read frequencies.

| SNP | Coverage | Variant read frequency | Sample | Verification |
|---|---|---|---|---|
| Mu_02 | 2,348 | 0.1 | M_08 | √ |
| | 2,386 | 0.1 | M_10 | √ |
| | 3,097 | 0.1 | M_12 | √ |
| Mu_03 | 834 | 0.08 | M_08 | √ |
| | 949 | 0.08 | M_10 | √ |
| | 1,162 | 0.06 | M_12 | √ |
| Mu_04 | 919 | 0.19 | M_10 | √ |
| Mu_05 | 726 | 0.43 | M_08 | √ |
| | 730 | 0.42 | M_10 | √ |
| | 916 | 0.41 | M_12 | √ |
| Mu_07 | 556 | 0.06 | M_12 | √ |
| Mu_09 | 1,055 | 0.05 | M_12 | × |
| Mu_10 | 966 | 0.06 | M_12 | × |
| Mu_11 | 34 | 0.29 | M_12 | √ |
| Mu_12 | 1,604 | 0.21 | M_12 | √ |

**Supplementary Table 8: Detailed information for 9 verified intra-module low-frequency SNPs.** They were verified in one to three modules, depending on the SNP calling results. The verification results can be found in Supplementary Figure 11 and Supplementary Table 7.

Gelöscht: 0

Gelöscht: 5

| Locus | Contig No. | Position | Enzyme | Restriction site | 5' Fluorescent dye | Forward primer | Reverse primer |
|---|---|---|---|---|---|---|---|
| Mu_02 | LFYR01000036.1 | 459,138 | HaeIII | GG^_CC | FAM | CAGGGTAACACACAACCTGGA | TGCTTTTTCTTCAACCTGTTCCT |
| Mu_03 | LFYR01000337.1 | 71,847 | HaeIII | GG^_CC | FAM | TGGAACACCGAATCTGCACA | GCGGAACACATAAAGGGGGA |
| Mu_04 | LFYR01000584.1 | 263,636 | HaeIII | GG^_CC | FAM | AATGCAAAGCCCCAACATCG | ATTTGAACCCATTGCGCTGG |
| Mu_05 | LFYR01000601.1 | 320,610 | HaeIII | GG^_CC | FAM | CGGTGCGAAAATTCAGGTGG | AGGACTTTGTGTGACAGACTTCT |
| Mu_07 | LFYR01000850.1 | 181,739 | HaeIII | GG^_CC | FAM | CCAGGGAGAGACAAGAACGAC | AGTGTTGCTGTTGCTGCATG |
| Mu_09 | LFYR01000907.1 | 145,314 | HaeIII | GG^_CC | FAM | CTTGCCTTCCCCATGACCAT | TTCTGTCACCAAATCGCCGA |
| Mu_10 | LFYR01000907.1 | 188,608 | HaeIII | GG^_CC | FAM | CTTGCCTTCCCCATGACCAT | GCCAGCTCGCGTCATTAATG |
| Mu_11 | LFYR01001228.1 | 68,748 | HaeIII | GG^_CC | FAM | CGATTCACACAGGCAATCGC | CGGCGACTGGACTTAGGTTT |
| Mu_12 | LFYR01001279.1 | 783,253 | HaeIII | GG^_CC | FAM | AGTGTGATGTTGGGTGAGACA | ACAGACTTGGACAGCATGGG |

**Supplementary Table 9: Significantly enriched GO terms for low-frequency SNPs and fixed SNPs, respectively.** The combination of all low-frequency (intra-module allele frequency <0.5) non-synonymous SNPs detected based on three 1000x-sequenced modules, and the combination of fixed non-synonymous SNPs detected based on twenty-four 81x-sequenced modules, were used for GO enrichment analysis.

| | | | | | |
|---|---|---|---|---|---|
| | **Low-frequency SNPs** | | | | |
| **BP (biological process)** | | | | | |
| GO.ID | Term | Annotated | Significant | Expected | Unjusted P |
| GO:0006468 | protein phosphorylation | 785 | 14 | 5.68 | 0.0013 |
| GO:0006075 | 1,3-beta-D-glucan biosynthetic process | 17 | 2 | 0.12 | 0.0065 |
| GO:0006422 | aspartyl-tRNA aminoacylation | 1 | 1 | 0.01 | 0.0072 |
| **CC (cellular component)** | | | | | |
| GO.ID | Term | Annotated | Significant | Expected | Unjusted P |
| GO:0098797 | plasma membrane protein complex | 35 | 3 | 0.19 | 0.00083 |
| GO:0005874 | microtubule | 21 | 2 | 0.11 | 0.00563 |
| **MF (molecular function)** | | | | | |
| GO.ID | Term | Annotated | Significant | Expected | Unjusted P |
| GO:0016165 | linoleate 13S-lipoxygenase activity | 5 | 2 | 0.04 | 0.00056 |
| GO:0004674 | protein serine/threonine kinase activity | 582 | 11 | 4.42 | 0.00463 |
| GO:0003843 | 1,3-beta-D-glucan synthase activity | 17 | 2 | 0.13 | 0.00721 |
| GO:0005200 | structural constituent of cytoskeleton | 17 | 2 | 0.13 | 0.00721 |
| GO:0004815 | aspartate-tRNA ligase activity | 1 | 1 | 0.01 | 0.00759 |
| GO:0035639 | purine ribonucleoside triphosphate binding | 2262 | 27 | 17.17 | 0.00933 |
| | **Fixed SNPs** | | | | |
| **BP (biological process)** | | | | | |
| GO.ID | Term | Annotated | Significant | Expected | Unjusted P |
| GO:0006464 | cellular protein modification process | 1011 | 35 | 16.81 | 0.00012 |
| **CC (cellular component)** | | | | | |
| GO.ID | Term | Annotated | Significant | Expected | Unjusted P |
| GO:0005634 | nucleus | 542 | 18 | 8.11 | 0.0015 |
| **MF (molecular function)** | | | | | |
| GO.ID | Term | Annotated | Significant | Expected | Unjusted P |
| GO:0005515 | protein binding | 3074 | 93 | 58.9 | 4.10E-06 |
| GO:0003830 | beta-1,4-mannosylglycoprotein 4-beta-N-acetylglucosaminyltransferase activity | 4 | 2 | 0.08 | 0.0021 |
| GO:0046914 | transition metal ion binding | 1161 | 36 | 22.25 | 0.0027 |
| GO:0015450 | P-P-bond-hydrolysis-driven protein transmembrane transporter activity | 7 | 2 | 0.13 | 0.0072 |
| GO:0008236 | serine-type peptidase activity | 148 | 8 | 2.84 | 0.0077 |

**Supplementary Table 10: Level of fixed SNPs in different species.**

| Species (genome size) | Sample source | Estimated age (years) | DNA source | Sequenced samples | Average mutation per sample | Pairwise-sample differentiation | Data source |
|---|---|---|---|---|---|---|---|
| *Prunus mira* (225 Mb) | one tree | 600 | Leaf | 32 | 12.7 | - | Wang et al. 2019 |
| | one tree | 550 | Leaf | 12 | 23.9 | - | |
| | one tree | 420 | Leaf | 23 | 17.7 | - | |
| | one tree | 300 | Leaf | 9 | 12.8 | - | |
| *Prunus persica* (225Mb) | one tree | 21 | Leaf | 23 | 3.74 | - | |
| | | | Root | 13 | 29.8 | - | |
| | one tree | 25 | Leaf | 16 | 6.19 | - | |
| | | | Petal | 13 | 11.31 | - | |
| | one tree | 30 | Leaf | 26 | 6.46 | - | |
| | one tree | 50 | Leaf | 8 | 6.25 | - | |
| | one tree | 40 | Leaf | 16 | 3.56 | - | |
| | one tree | 2 | Leaf | 75 | 1.97 | - | |
| *Prunus mune* (220 Mb) | one tree | 20 | Leaf | 25 | 12.9 | - | |
| | | | Root | 32 | 25.4 | - | |
| | one tree | 8 | Leaf | 33 | 5.7 | - | |
| *Salix suchowensis* (480 Mb) | one tree | 1 | Leaf | 19 | 1.26 | - | |
| | | | Root | 21 | 2.86 | - | |
| *Brachypodium distachyon* (272 Mb) | one plant | 1 | Leaf | 29 | 3.17 | - | |
| | | | Root | 8 | 4.75 | - | |
| | | | Lemma | 7 | 2.57 | - | |
| *Fragaria vesca* (210 Mb) | one plant | 1 | Leaf | 45 | 1.93 | - | |
| | | | Stem | 4 | 4.75 | - | |
| *Arabidopsis thaliana* (119 Mb) | two plants | 1 | Leaf | 64 | 0.69 | - | |
| *Oryza sativa* (373 Mb) | four plants | 1 | Leaf (Tiller) | 29 | 4.79 | - | |
| | | | Leaf (Callus) | 13 | 194.8 | - | |
| *Quercus robur* (720 Mb) | one tree | 234 | Leaf | 2 | - | 38-47 | Schmid-Siegert et al. 2017 |
| *Zostera marina* (204 Mb) | one genet | 750 – 1,500 | Meristematic region and the basal portions of the leaves | 24 | - | 1,216 (on average) | Present study |

Schmid-Siegert, Emanuel, et al. "Low number of fixed somatic mutations in a long-lived oak tree." Nature plants 3.12 (2017): 926.

Wang, Long, et al. "The architecture of intra-organism mutation rate variation in plants." PLoS biology 17.4 (2019): e3000191.
